# Rapidly evolving aphid gall effector proteins exhibit saposin-like folds

**DOI:** 10.64898/2026.03.27.712717

**Authors:** Fatema Bhinderwala, Aishwarya Korgaonkar, Kota Gopalakrishna, Thomas C. Mathers, Shuji Shigenobu, J. Fernando Bazan, Saskia A. Hogenhout, Guillermo Calero, Angela M. Gronenborn, David L. Stern

## Abstract

Many insects manipulate plants by injecting effector proteins. In one extreme example of this molecular “hijacking”, *Hormaphis cornu* aphids inject bicycle proteins into *Hamamelis virginiana*, contributing to the development of novel organs called galls. Bicycle proteins share no amino acid sequence similarity with proteins of known function. Here, we report the crystal structures of two divergent bicycle proteins. Both proteins contain saposin-like folds: one with multiple disulfide bonds exhibits a swapped domain topology; the other has no disulfide bonds and possesses two distinct, tandem domains. To explore the structural evolution of bicycle proteins, we attempted to predict bicycle protein structures with Alphafold2 (AF2) and other deep learning programs. While AF2 did not recover the two experimental structures using existing databases, it succeeded when provided with multiple sequence alignments (MSAs) of protein sequences from newly sequenced closely related species. Using this approach, we generated 2400 high-confidence bicycle protein predictions from seven aphid species. While all aphid bicycle proteins contain predicted saposin-like folds, they display a vast diversity of structural and physicochemical properties. While this diversity thwarts prediction of conserved functions encoded in structure, it suggests that bicycle proteins have evolved to target diverse plant processes and/or to evade plant immune surveillance. Our extension of AF2 with custom MSAs of proteins from closely related species provides a generalizable, powerful approach for predicting structures of rapidly evolving protein families.

**Significance statement:** Parasites introduce specialized “effector” proteins into hosts to suppress host immunity and to release nutrients. The molecular functions and structures of most effector proteins are unknown. Effector proteins often evolve rapidly and share no similarity with proteins of known function. Here, we demonstrate that machine learning algorithms can predict the structures of aphid “bicycle” effector proteins when supplemented with data from closely related species. We exploit this finding to generate predictions of 2400 bicycle protein structures. Aphid bicycle proteins exploit a common folding motif, yet exhibit topologically distinct structures that form separate structural clusters. Despite the clustering of these proteins in structure space, they occupy a nearly uniformly physicochemical space, suggesting that they encode a large diversity of molecular functions.

## Introduction

Pathogens employ effector proteins to manipulate host physiology, development, and behavior (1–6). In response to these effectors, hosts deploy immune responses. Over evolutionary time, conflict between effectors and immune responses can result in a so-called molecular “arms-race” between the host and pathogen, associated with rapid molecular evolution (7–11). This results in the generation of novel protein families that share little amino acid sequence similarity with proteins of known function, complicating the inference of their molecular functions from sequence-based comparisons.

Aphids are plant sap-sucking insects that secrete salivary proteins into plant cells to facilitate access to nutrients and inhibit plant defense responses (12). The most extreme examples of plant manipulation by aphids and other insects is the production of novel organs, called galls, which reflect reprogramming of plant cell growth and development (13, 14). Aphids and other insects induce galls to provide nutritious enclosed spaces that protect the insects from predators, parasites, and environmental vicissitudes (15–17). Bicycle proteins from *Hormaphis cornu* have recently been implicated as aphid-produced effectors that influence gall development (13). In particular, the bicycle protein Determinant of Gall Color specifically regulates a single anthocyanin biosynthetic pathway by suppressing pigment-related gene expression to turn red galls green (13). It is not currently clear how individual bicycle proteins perform such potent and specific manipulations of plant biology. *Bicycle* genes have evolved rapidly owing to strong diversifying selection (13), and the genome of the gall-inducing aphid *H. cornu* encodes more than 650 *bicycle* paralogs. All aphid genomes examined to date encode *bicycle* genes (18), suggesting that many or all aphids inject bicycle proteins into plants to manipulate plant physiology and development.

Bicycle proteins possess an N-terminal secretion signal sequence, which is likely removed during translocation to the secretory vesicles. A pair of distributed cysteine-tyrosine-cysteine motifs (C-Y-C) is found in most bicycle proteins—giving the family its name (“bi-CYC-like”)—and suggests that these proteins contain two structurally related domains. However, outside of the signal sequences, these proteins display no similarity to proteins or domains of known function (13). Here, we report the X-ray structures of two bicycle proteins from *H. cornu*. Both structures contain saposin-like folds, but in topologically different arrangements. These experimentally determined structures were used as positive controls to benchmark AlphaFold2 (AF2) predictions of additional bicycle proteins. Multiple sequence alignments (MSA) of bicycle proteins encoded by newly sequenced genomes of species closely related to *H. cornu* allowed AF2 to predict models that closely resembled the crystal structures, as well as high-confidence models for thousands of bicycle proteins from multiple species. These predicted models exhibit extensive variability in overall amino acid sequence space, especially of surface residues, as well as of physicochemical properties, with no evidence of conserved surface accessible domains that might define biochemical functions. The extensive diversity of these proteins may be required to allow them to engage with multiple molecular targets in plants and/or to evade plant defense mechanisms.

## Results

### A bicycle protein with helix swapped saposin-like folds

We selected 18 *bicycle* genes (Table S1) that are highly expressed in *H. cornu* salivary glands based on RNAseq data (13) for heterologous expression. Of these, four proteins expressed well in *E. coli* and three in *SF9* insect cells and were purified in sufficient quantities for initial crystallization tests. Of these, two proteins—g3873 and g2703—yielded diffraction-quality crystals (Figure S1, Table S1).

The X-ray structure of g3873 was solved at 2.0 Å using sulfur-SAD phasing and revealed an all-helical fold, possessing two sub-domains, each comprising four-helical bundles: helices □1, □2, □3, and □4’ form one bundle, and □1’, □2’, □3’, and □4 form the second bundle (Figure 1A,C, and S2, Table S2). The first (C30) and second (C191) conserved cysteines of the first and second CYC motif, respectively, form a disulfide bond linking helices □1 and □4’. The second (C92) and first (C120) conserved cysteines of the first and second CYC motif, respectively, form a second disulfide bond linking helices □4 and □1’ (Figure 1A). A third disulfide bond connects C38 in helix □1 and C48 in helix □2, and these cysteines are not strongly conserved across bicycle proteins. The average distance between the sulfur atoms in the C30-C191, C92-C120, and C38-C48 bonds is 2.05 ± 0.1 Å, 2.08 ± 0.1 Å, and 2.03 ± 0.05 Å, respectively. The presence of three disulfide bonds as well as an additional free sulfhydryl group was confirmed by titration using 5,5⍰-dithiobis-2-nitrobenzoic acid (19, 20) and the sulfur positions were clearly identified in the difference density maps (Figure S3). The two disulfide bonds associated with the CYC motifs form left-handed (LH) spirals, with χ1 and χ1⍰ dihedral angles of 180° and 60°, respectively. The C30-C191 disulfide bond is also an LH spiral, whereas the C92-C120 disulfide bond is a right-handed (RH) staple. Of the 20 different disulfide bond geometries, -LH spirals are the most common and energetically favorable disulfide conformation, with the lowest dihedral strain energy (DSE) (21). The calculated DSE is 9.2±1.1 kJ/mol for each of the two LH spirals and 14.9±1.2 kJ/mol for the RH staple (Table S3) (22, 23).

**Figure 1:**
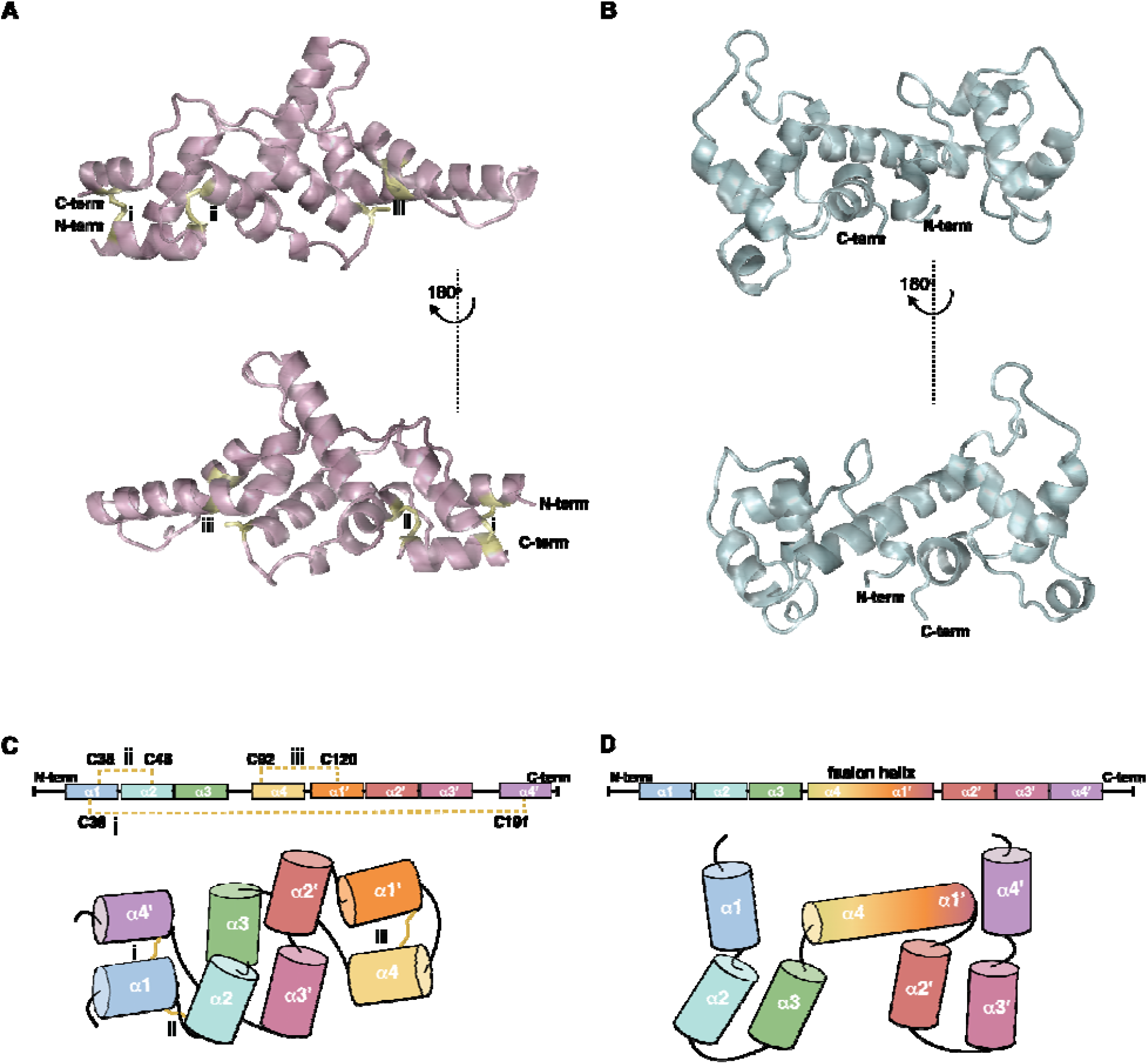
X-ray structures of two *H. cornu* bicycle proteins. (A) 2.0 Å X-ray structure of G3873 in ribbon representation (pink) with the three disulfide bonds and the free cysteine residue shown in stick representation (yellow), embedded in a gray surface representation. (B) 1.4 Å X-ray structure of G2703 in ribbon representation (aqua) embedded in a gray surface representation. (C and D) Schematic representation of the secondary structure elements in the G3873 (C) and G2703 (D) polypeptide chains. Helical elements are represented by colored boxes or cylinders, labeled □1 through □4’. In G3873, the left and right side four-helical bundles include □1, □2, □3, and □4’ and □1’, □2’, □3’, and □4 helices, respectively. In G2703, the left- and right-side four-helical bundles include □1, □2, □3, and the □4 portions of the □4-□1’ fused long central helix, and the □1’ portion of the fused long central helix, □2’, □3’, and □4’ helices, respectively. Color coding runs from the N-terminus (aqua blue) to the C-terminus (purple). Disulfide bonds are indicated by yellow lines (dashed; top; zig-zag bottom)

The structure of g3873 (Figure 1A, C) is dissimilar from all proteins in the PDB (24), as determined by DALI (Z-score>8) (25) and FoldSeek (E-value threshold = 0.001, sensitivity = 9.5) (26) (Table S4, Supplementary Methods). Manual inspection revealed, however, that each set of four clustered alpha helices is reminiscent of the saposin fold (27– 29). Saposins are conserved proteins that contain a hydrophobic core sandwiched between two pairs of alpha helices, which allows them to bind phospholipids (30). Although the two four-helical bundles of G3873 resemble the canonical saposin fold, they appear to lack a hydrophobic internal surface that can adopt an alternate open conformation (31).

### A bicycle protein with tandem saposin-like folds

The second bicycle protein we crystallized, g2703, has a highly divergent amino acid sequence from other bicycle proteins, with no cysteines, but it has a similar intron-exon structure to other *bicycle* genes and was identified as a member of the bicycle gene family using a gene structure classifier (18) (Figure S4). This protein has an N-terminal secretion signal and is highly expressed in the salivary glands of *H. cornu* (Figure S5), supporting the inference that it serves as an effector protein. Thus, while *bicycle* genes were named originally based on the presence of conserved C-Y-C motifs, the sequence of g2703 suggests that at least some *bicycle* genes do not fit neatly within this sequence-based categorization.

The X-ray structure of the g2703 protein (Figure 1B, D) was solved at 1.4 Å using selenomethionine-labeled protein (Figure S1) *and multiwavelength anomalous diffraction (MAD)* phasing. The structure of g2703 contains two saposin-like domains in tandem (Figure 1D), connected by a long alpha helix. This long central helix appears to have evolved by a merger of the last helix of the first saposin-like unit and the first helix of the second unit (Figure S6). Similar to g3873, neither of the two saposin-like folds exhibits strong amphipathic properties that might facilitate protein-lipid interaction (32).

The saposin-like domains of g3873 and g2703 share limited structural similarity (33, 34) with the 52 saposin structures deposited in the PDB (24) (Figure S7 and S8, Table S5 and S6). Most similar to g3873 is the saposin domain of a proteolytically processed human acyloxyacyl hydroxylase saposin (35) (PDB ID: 5W78, TM_score = 0.44 Å, backbone RMSD for 68 residues = 3.65Å) (Figure S7). The N-terminal saposin-like domain of g2703 showed modest similarity to the saposin domain of prophytepsin (36) from barley, characterized as a vacuolar aspartic proteinase (PDB ID: 1QDM, TM_score = 0.495, backbone RMSD for 71 residues = 4.03 Å) (Figure S9).

The presence of saposin-like domains in the crystal structures of both g3873 and g2703 suggests that they evolved from a common ancestral protein, despite the absence of significant sequence similarity between these two proteins. This observation provides independent support for the hypothesis that the bicycle gene-structure classifier (18) can identify *bicycle* homologs that cannot be recognized by sequence similarity.

### AlphaFold2 can predict bicycle protein structures using custom multiple sequence alignments

Since these two bicycle proteins exhibited no significant sequence similarity to others in Genbank (37) and the solved structures were different from all structures in the PDB (24), we submitted our structures as targets for the 15th Critical Assessment of Techniques for Protein Structure Prediction (CASP15) challenge (38, 39). CASP is a community-wide experiment held every two years that assesses computational methods for determining protein structures from amino acid sequences. At the CASP15 competition, in the single protein and domain-modeling category, 162 groups submitted predictions for g3873 (CASP ID: T1130) and g2703 (CASP ID: T1131). Final metric GDT_TS scores were used to evaluate the accuracy of the predictions relative to the experimental structures (40). As expected, given the breakthrough success of AlphaFold2 (AF2) in CASP14 (41, 42), most prediction teams deployed deep learning-based algorithms based on AF2, resulting in >90 GDT_TS scores for most targets. The two groups that accurately predicted the structure of g3873 (T1130) (GDT_TS scores >50) in CASP15 incorporated MSAs of bicycle proteins reported in one of our earlier studies (18). In contrast, no group correctly predicted g2703 (mean GDT_TS score of 12, range 11.65-25.31), making g2703 (CASP ID: T1131) the lowest-scoring target (43). This may be because homologs of g2703 were not present in any previously released genome sequences and thus could not be used to generate MSAs. Consistent with the results from CASP15, we also failed to generate accurate predictions for both g2703 or g3873 using AF2, AF3, and ESM-Fold in-house with publicly available sequence databases (44) (Figure 2A and C, S10).

**Figure 2:**
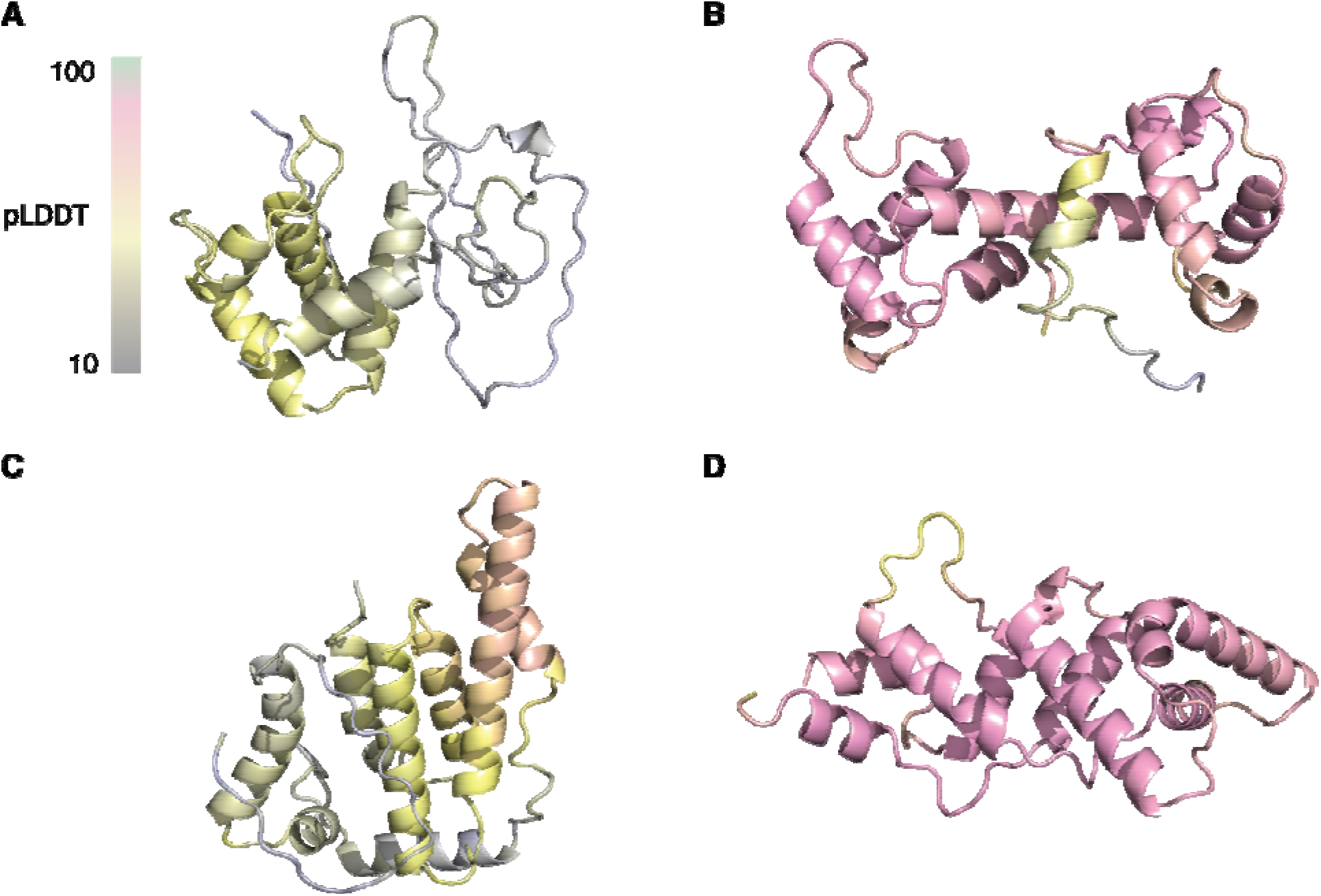
AF2 predicted models for G2703 (A and B) and G3873 (C and D). In (A) and (C), no MSAs were used because the default databases used by AF2 for homolog searches do not contain any homologs. In (B) and (D) custom MSAs were generated with homologous proteins from closely related species. The AF2 predicted models are colored by per-residue measures of local confidence (bar) in the predicted local distance difference test (pLDDT). Grey and pink represent ∼10 and ∼85 confidence levels, respectively.

Given these observations, we hypothesized that MSAs from homologous proteins are required for accurate reconstruction using current machine learning algorithms for structure prediction. We therefore sequenced and annotated the genomes of three species closely related to *H. cornu*: *H. betulae, H. hamamelidis*, and *Hamamelistes spinosus* (Figure S11A). In addition, we re-annotated genomes from several aphid species distantly related to *H. cornu* (13), *Schlechtendalia chinensis* (45), *Tetraneura akinire* (18), and *Acyrthosiphon pisum* (46), guided by publicly available RNA-seq data (www.ncbi.nlm.nih.gov/sra). We predicted *bicycle* genes for all species using a gene-structure classifier (18). Strikingly, we observed 1044 and 800 *bicycle* genes in *H. spinosus* and *S. chinensis*, respectively, revealing very large numbers of *bicycle* genes in these gall forming species and greatly increasing the number of predicted bicycle proteins that were available for study. Using previously published RNA-seq data for *S. chinensis*, we found that *bicycle* genes were expressed at highest levels in the gall foundress generation (Figure S11B), consistent with the hypothesis that bicycle proteins contribute to gall development in this species, as they do in *H. cornu* (13).

Searching the translated aphid proteomes from these seven species with *phmmer* (47) identified 197 and 11 homologs of g3873 and g2703, respectively. Despite the fact that bicycle proteins were not part of the training set used to parameterize AF2, AF2 accurately predicted the experimentally determined X-ray structures of both proteins when provided with these custom MSAs (Figure 2B, 2D, and S12).

Given the accuracy of the AF2 predictions for the two experimentally solved bicycle proteins, we explored whether AF2 could generate high confidence predictions for 4924 bicycle proteins from seven aphid species (Figure S11C-F, Table S7). MSA depth was positively correlated with prediction confidence (Figure S11F) and AF2 predicted 2400 high confidence bicycle protein structures (≥80% of amino acids with pLDDT ≥ 60 over at least 70 residues) .

As discussed below, aphid bicycle proteins are predicted to occupy a diverse structural space, so it is unclear whether benchmarking with two X-Ray structures provides strong confidence for the full range of predictions. However, several observations suggest that these AF2 predicted structures provide a plausible snapshot of the true diversity of bicycle protein structures. First, we benchmarked AF2 on bicycle proteins with two extremely different experimentally derived structures (g3873 and g2703), providing some confidence that AF2 can predict diverse bicycle protein structure. Second, as discussed further below, all of the high confidence predictions include saposin-like folds. And, third, we focus only on predictions with the high average pLDDT scores, which has been shown to be a reliable measure of fold accuracy (48).

To explore the robustness of this approach, we examined the effect of MSA depth on AF2 performance. First, we predicted structures using randomly resampled MSAs of g3873 and g2703 (Figure S13A-D). The MSA for g3873 was more informative because of the greater number of homologs available for this protein. Predictions with MSAs of at least 16 homologs generated predictions with similar accuracy to the full MSA of 197 homologs (Figure S13A-B). The analysis of G2703 showed a similar trend (Figure S13C-D). To determine if this result generalized, we randomly subsampled MSAs from 100 randomly chosen bicycle proteins with high confidence structure predictions and again observed that accuracy leveled off for predictions using MSAs with at least 16 homologs (Figure S13E-F). Thus, sequencing genomes of clusters of closely related species provides a potentially powerful method to generate informative MSAs for protein structure prediction of rapidly evolving protein families.

### Bicycle proteins exhibit extensive structural diversity and surface variability

We used this dataset of 2400 high confidence bicycle protein predictions to explore four questions: Is the saposin-like fold conserved across aphid bicycle protein structures? Are the tandem and helix-swapped permuted structures related? How conserved is the bicycle protein structural and physicochemical space? Finally, does the conformational space and the associated physicochemical signature of bicycle proteins permit inference of a putative function for bicycle proteins?

#### Saposin-like folds are found in all aphid bicycle proteins

All high-confidence predicted folds possess at least one saposin-like domain. Nonetheless, these domains could not be perfectly superimposed and instead displayed a wide variety of quantitative variations in saposin-like domain shape, size, and arrangement, as we discuss in more detail below.

#### The tandem-linked bicycle proteins likely evolved from helix-swapped proteins

Almost all of the bicycle proteins in our database encode proteins with helix-swapped saposin-like domains. We therefore wondered how the tandem-linked bicycle proteins, like g2703, evolved. *Bicycle* genes are often found in large paralog groups, apparently reflecting historical gene duplication and divergence (13). We therefore examined the genomic context near *g2703* and identified four putative paralogs. The proteins encoded by these paralogs share almost no sequence similarity to g2703 (Figure S4, S6), putting them in the “twilight zone” of protein similarity (49). Strikingly, these four putative paralogs are predicted to encode proteins with tandem-linked saposin-like domains, despite the predicted presence of disulfide bonds (Figure S4, S6). One plausible evolutionary model is, therefore, that most genes in this paralog group first evolved a tandem-linked arrangement with disulfide bonds and g2703 evolved by duplication from these paralogs and subsequent loss of cysteines and most recognizable sequence similarity.

Mapping protein structure and intron-exon structure onto a multiple sequence alignment of these paralogs revealed that the alpha-helical “handle” that connects the two tandem-linked saposin-like domains of g2703 likely evolved through fusion of the last and first helices from the first and second saposin-like domains, respectively (Figure S6). This multiple sequence alignment also shows gene intron positions and reveals the extreme diversity of exon number and protein sequence, even between these genes from a single paralog family. It is striking that *g2703* is one of the most highly expressed *bicycle* genes in *H. cornu* salivary glands (Figure S5), implying that despite its relatively recent origin and highly divergent sequence, it has acquired an important function for aphid biology during gall development.

#### Bicycle proteins exhibit diverse sequences and structures

Given this evidence that bicycle proteins can evolve diverse structures, we were motivated to further explore bicycle protein diversity. We first characterized protein sequence divergence using pairwise length-normalized Levenshtein distances (50) (lower half below diagonal of Figure 3A). Almost all sequences are highly divergent from each other, with most pairwise distances > 0.75. The dendrogram adjacent to the distance matrix (Figure 3A) reveals that some sequences cluster together within species, especially for *S. chinensis*, but even these sequences do not exhibit clear clusters of similarity that might define subfamilies. Thus, bicycle proteins display little evidence of sequence similarity, extending our initial observations of the extreme diversity of bicycle protein sequences within species (13, 18).

**Figure 3:**
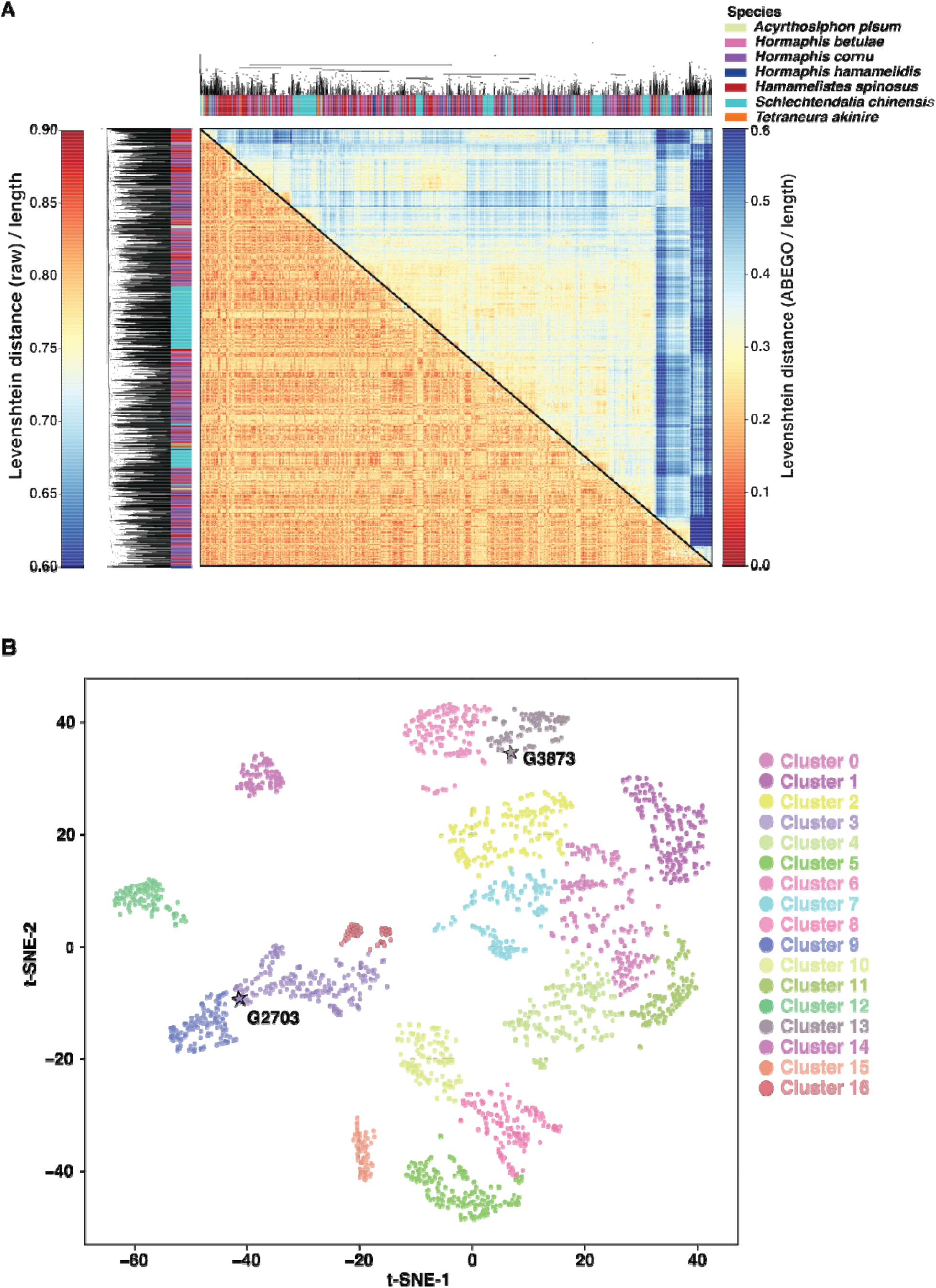
Bicycle proteins are predicted to adopt a wide diversity of structures. (A) Heatmap of pairwise normalized Levenshtein distances for 2400 bicycle proteins from seven species computed using amino-acid sequences (lower) and ABEGO-encoded backbone-state strings derived from each AF2-predicted model (upper). In both heatmaps, colors represent distance (0 = identical strings; 1 = maximally different, normalized to the length). Bicycle amino acid sequences display extreme diversity (heatmap is colored blue to red, ranging from 0.6 to 1.0), even within species, whereas ABEGO strings retain intermediate similarity (heatmap distances colored from blue to red from 0.0 to 0.6), even across species. These patterns suggest that bicycle proteins exhibit some conserved higher-order backbone structural grammar despite low primary sequence identity. The dendrograms were calculated by hierarchical clustering the distance measures shown in the heatmaps. Dendrogram tips are colored by species (see key), revealing that amino acid sequences and protein structures are divergent across all aphid species sampled. (B) t-SNE embedding of all-vs-all TM scores for 2400 bicycle proteins from seven species. Each point represents one protein and is colored according to its cluster assignment (k_cluster_ = 17). Structurally similar proteins (higher TM scores) are located close to each other.

We then explored AF2 predicted structural diversity by calculating Levenshtein distances using the ABEGO classification (51) of structural motifs of amino acids (upper half of Figure 3A). The ABEGO-based map exhibits a wider dynamic range than the raw amino-acid sequence-based map, with distances saturating at approximately 0.5 rather than near 0.8 (Figure 3A), suggesting that bicycle proteins do share similarities in ABEGO space. Also, the ABEGO distance matrix displays some clustering, although little of this clustering is by species (dendrogram on top of Figure 3A).

To further explore the clustering observed in the ABEGO-based map, we performed t-SNE (52) embedding of all-vs-all TM-align comparisons for all 2400 proteins (Figure 3B). Many distinct structural neighborhoods were identified in this embedding. Medoid models for each cluster illustrate that bicycle proteins occur in a wide variety of arrangements, including one, two, three, four, or six saposin-like domains (Figure 4). Substantial species partitioning was observed for some clusters (Figure S14, S15). This is most striking for the phylogenetically isolated species in our study: *A. pisum, T. akinire*, and *S. chinensis*. Nonetheless, bicycle proteins from *Hormaphis* and *Hamamelistes* occupy most of these clusters, indicating that the relative arrangements of the alpha helices in the saposin-like domains vary extensively, even within species. Thus, the bicycle protein family has evolved to occupy a large and discontinuous region of structural space within the context of the saposin-like four-helix domains.

**Figure 4:**
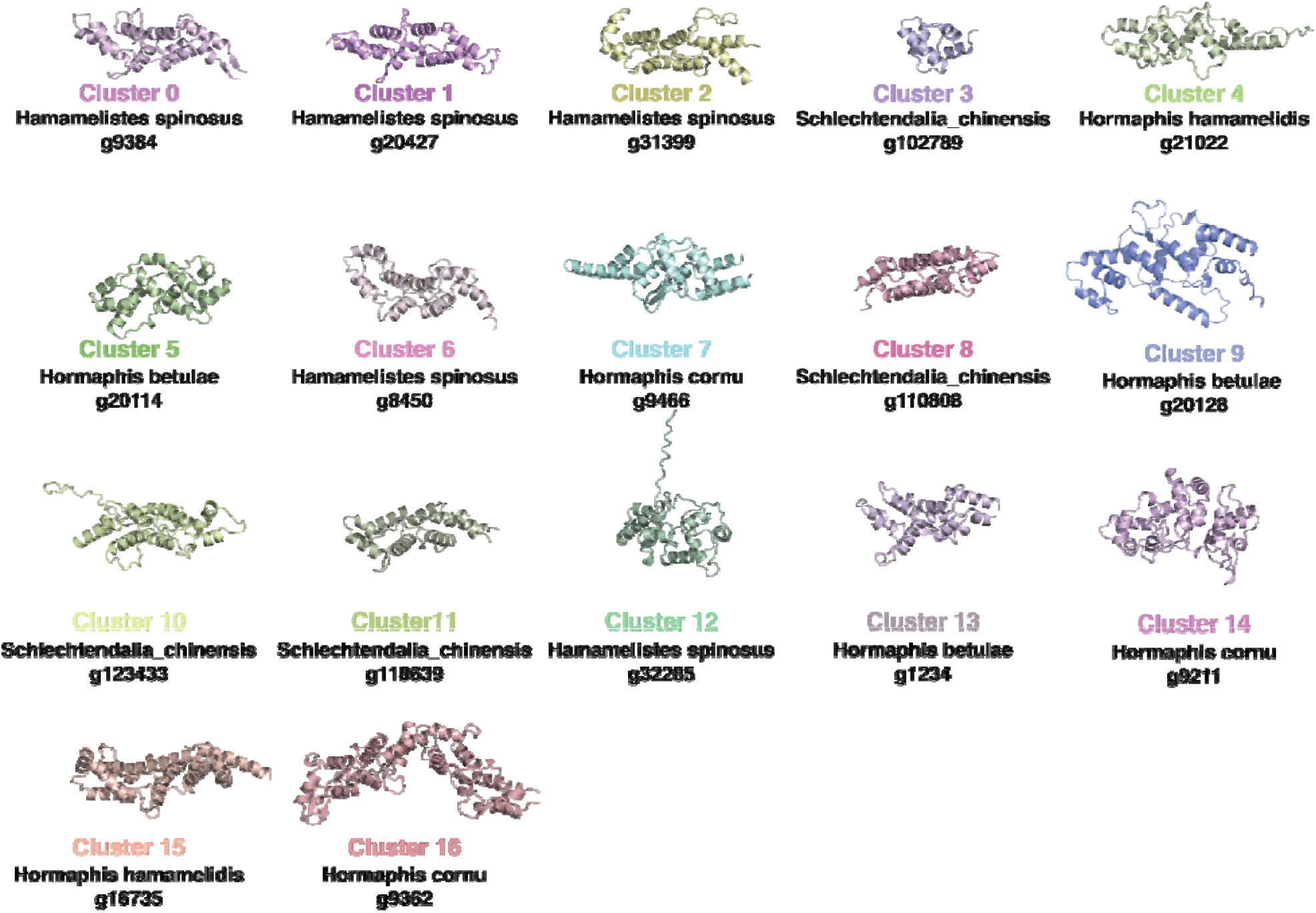
Ribbon models of the medoids representative for each Leiden cluster from Figure 4B shown in their corresponding cluster colors. The medoid possesses the highest average TM-score (the lowest average distance) to all other members of that cluster.

#### Bicycle proteins occupy diverse physicochemical spaces

In an effort to identify features that may be conserved among the bicycle proteins, we assessed the physicochemical features of each predicted protein. Five complementary properties were examined: surface chemistry, electrostatic charge, amphipathic character, surface hydrophobic patchiness, and surface texture (Tables S7 and S8). We embedded these features using UMAP (52) and attempted to identify groups of similar proteins using Leiden clustering (50) (Figure 5). Even though UMAP tends to “aggressively” cluster points (53, 54), we observe an almost uniform distribution of bicycle proteins characterized by these five properties, suggesting that bicycle proteins sample relatively uniformly across this physicochemical space.

**Figure 5:**
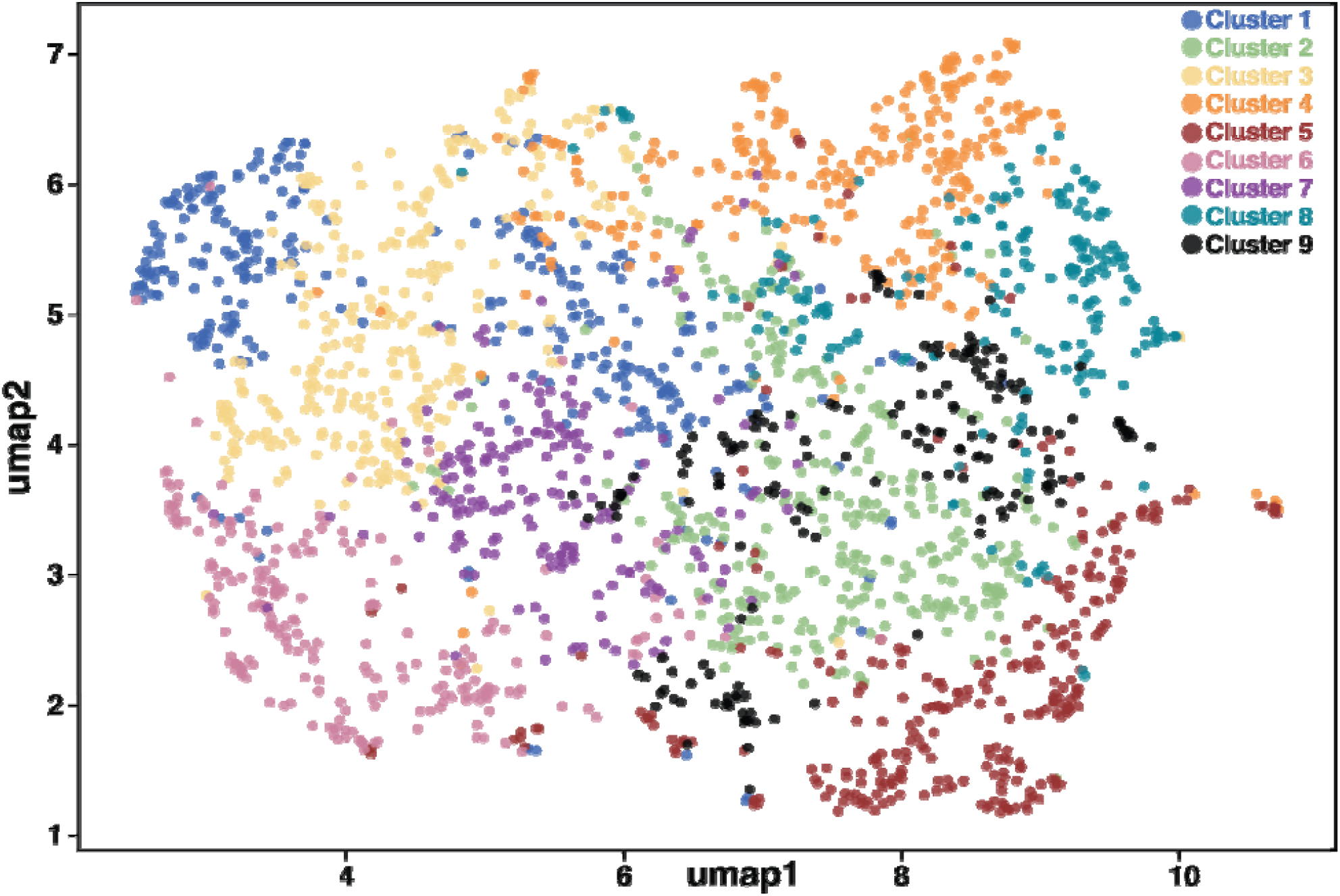
Bicycle proteins display a wide and almost continuous diversity of physicochemical features in the first two dimensions of a UMAP embedding. Each point represents one protein. Different Leiden clusters are shown in different colors and contain sets of proteins primarily with different quantitative combinations of surface hydrophobicity, charge balance, and amphipathicity

We observed only a few interpretable subgroups in this physicochemical space, defined primarily by distinct combinations of surface hydrophobicity, charge balance, and amphipathicity. One subset of clusters is defined primarily by hydrophobicity and patch morphology: clusters 1, 6, and 3 combine hydrophobicity with minimal amphipathicity. A second subset was dominated by electrostatics and charge composition, being either positively charged (cluster 2), negatively charged (cluster 4), or possessing a pronounced surface dipole (cluster 5). The third subset possesses amphipathic helices and high hydrophobic moment (clusters 7 and 9) or low hydrophobicity with amphipathic signatures (cluster 8). Poisson-Boltzmann surface electrostatics were superimposed on the mediod structures of each cluster to aid visualization of these physicochemical subtypes (Figure S16). The physicochemical subtypes are not species-partitioned (Figure S17, S18, and S19), suggesting that the variability in physicochemical features extends across the bicycle protein family.

#### Lack of conserved domains thwarts the prediction of bicycle protein functions

The prior analysis suggests that there is limited conservation of features that might provide clues about the molecular function of bicycle proteins. Therefore, we examined whether surface residues displayed any patterns of conservation. We mapped Shannon entropy of residues derived from a multiple sequence alignment of bicycle proteins, which provides a quantitative measure of per-residue conservation, onto the crystal structure of g3873 (Figure S20). All conserved residues are buried within the interior of the protein, likely required for structural packing, whereas all surface residues show extreme variability.

To assess whether bicycle proteins share structural similarities with known cysteine-rich insect protein families, the disulfide connectivity of all predicted bicycle protein models from seven aphid species was classified (Figure S21). Three dominant Cys-connectivity patterns were identified connecting C1-C4, C2-C3; C1-C6, C2-C3, C4-C5; and 1C-C5, C3-C4 with a free C2), together accounting for ∼28% of the dataset. None match the interlocking C1-C3, C2-C5, C4-C6 topology of classic odorant-binding proteins (OBPs) (55), nor the Plus-C or Atypical variants, ruling out OBP family membership.

To examine the energetic landscape underlying this conservation pattern, MSA-level frustration analysis (FrustraEvo) was performed on the three dominant topology subgroups (Figure S22). The conserved energetic scaffold consists exclusively of the disulfide staples and a small hydrophobic or aromatic core, with all other positions energetically neutral. Per-residue frustration versus solvent accessibility further showed that maximally frustrated residues are approximately threefold enriched in the buried decile (∼60%) compared with the solvent-exposed decile, opposite to the surface-frustration re-patterning reported for fungal orphan candidate effectors (56). Surface residues remain predominantly neutral, consistent with the variability observed in the sequence entropy analysis. This absence of any specific conserved sequence signature on the exterior of the protein precludes an inference of bicycle protein functions from these data.

## Discussion

Pathogens frequently introduce effector proteins into hosts for multiple purposes, including driving physiological and developmental programs beneficial to the parasite, securing access to nutrients, and suppressing the host immune response (57, 58). These proteins often alter host cell structure, gene expression, or normal signaling pathways. Many effector proteins do not exhibit domains of known function, and the putative functions of most effector proteins are challenging to decipher (12, 59, 60). Protein structure can provide one path toward insight into effector functions.

Here we report the crystal structures of two *H. cornu* aphid effector proteins that belong to the bicycle protein family. The CYC motifs present in the g3873 structure contribute to two saposin-like folds, in a helix swapped arrangement, secured by disulfide linkages. In contrast, the g2703 structure includes tandem-linked saposin-like folds. It is striking that although g2703 includes no cysteines, and thus no CYC motifs, it also generates saposin-like folds. This shared structural motif suggests that g3873 and g2703 shared a common ancestor that contained saposin-like folds.

Despite the presence of saposin-like folds in these proteins, current evidence suggests that bicycle proteins do not possess functions similar to canonical saposins. Saposins are found across eukaryotes and participate in membrane recycling (27, 31). These proteins contain a strongly hydrophobic core that can be exposed to capture lipids. The structures of the bicycle proteins solved in this study do not display features that would allow a hydrophobic core to be exposed. Instead, the saposin-like fold apparently provides a rigid backbone for a rapidly evolving solvent-exposed surface. Other proteins without sequence similarity to saposins contain domain-swapped saposin-like folds (61), although in this case, pairs of helices are swapped, which is different from the single helix swap observed in g3873. Saposin-like folds have also been observed for other proteins, distinct from those that possess lipid or membrane-modifying properties (62, 63). Thus, the saposin-like fold is likely an example of convergent evolution in multiple independent protein families and may provide a stable backbone that allows diversification in sequence space for multiple functions (63). Another example of this kind of convergent evolution is found in the lipocone family of proteins that act in several functional contexts, including regulation of membrane lipid composition, extracellular polysaccharide biosynthesis, and biogenesis of lipopolysaccharides (64).

While AF2 and other deep learning and protein language models failed to accurately predict bicycle protein structures with default settings and existing databases, we found that AF2 can accurately predict these structures when provided with MSAs of homologous proteins identified from the genomes of closely related species. By sequencing several new aphid genomes and re-annotating previously available genomes to optimize the annotation of *bicycle* genes, we generated 2,400 high-confidence bicycle protein structures across seven aphids species. This large dataset allowed us to explore aphid bicycle protein structure evolution and to search for sequence, structural, and physicochemical patterns that may provide clues about bicycle protein function.

We observed high levels of sequence divergence amongst the bicycle proteins, (Figure 3A) with no clear clusters of similar sequences, even within a single species. In contrast, the predicted structure space, either ABEGO encoded (Figure 3A) or measured by structural similarity with TM scores (Figure 3B), displayed clusters of similar backbone structures (Figure 4). Individual clusters were mostly not associated with individual species (Figure S14 & 15), indicating that a broad diversity of protein shapes are encoded in bicycle proteins from multiple species.

In contrast to the clusters in shape space, we observed a nearly continuous distribution of bicycle proteins in physicochemical space (Figure 5), which is also not strongly partitioned by species. This variability is consistent with the lack of sequence conservation, particularly on the surface of bicycle proteins (Figure S20). There are thus no surface regions of these proteins that exhibit conservation in sequence or physicochemical space, even amongst subsets of proteins, that might guide inference of the molecular functions of these proteins. It remains unclear how proteins that exhibit such a remarkable diversity of structures and apparent physicochemical properties can accomplish potent and precise modifications of plant development (13).

The proteins within sub-clusters of the structural space (Figure 3B) do not tend to cluster within the physicochemical property groups shown in Figure 5, indicating that structure and physicochemical properties have evolved independently within this protein family. As a result, the diversity of protein structures and surface physicochemical properties highlight the challenge of inferring effector protein function strictly from structural data. Nonetheless, the presence of structural and amino acid sequence diversity exhibited by bicycle proteins, combined with our previous finding that *bicycle* genes have evolved in response to strong positive natural selection (13), suggests that individual bicycle proteins may perform distinct functions in plant cells (13) and/or that they have evolved to evade plant defense recognition systems.

Families of rapidly evolving protein families often participate in host-parasite interactions and inferring their structures and functions remains a challenging problem. We sequenced genomes of closely related species to generate informative MSAs, which enabled accurate protein structure prediction. The success of this approach suggests that large sequencing efforts should balance breadth of taxonomic sampling with focused sampling of multiple closely related species to facilitate large-scale prediction of protein structure for rapidly evolving protein families.

## Acknowledgements

We thank Goran Ceric, Robert Lines, Donald Olbris, and Tom Dolafi for assistance in installing, running, and maintaining the software and databases used in this study. For help generating the *Hormaphis betulae* genome, we thank Dr. Katsushi Yamaguchi for next-generation sequencing and bioinformatics support, Dr. Shunta Yorimoto for aphid collection, and Dr. Asahi Akita for next-generation sequencing support, including genomic DNA extraction and library preparation. We thank members of David Clapham’s lab for use of their Crystal Gryphon instrument and Teresa Brosenitsch for critical reading and editing of the manuscript. We acknowledge use of the Stanford Synchrotron Radiation Lightsource, SLAC National Accelerator Laboratory, which is supported by the U.S. Department of Energy, Office of Science, Office of Basic Energy Sciences under Contract No. DE-AC02-76SF00515. The SSRL Structural Molecular Biology Program is supported by the DOE Office of Biological and Environmental Research and by the National Institutes of Health (NIH Grant P30GM133894). This article is subject to HHMI’s Open Access to Publications policy. HHMI lab heads have previously granted a nonexclusive CC BY 4.0 license to the public and a sub-licensable license to HHMI in their research articles. Pursuant to those licenses, the author-accepted manuscript of this article can be made freely available under a CC BY 4.0 license immediately upon publication.

## Author contributions

Protein expression and crystallization was performed by KG, AK, and FBhinderwala. X-ray structures were solved by FBhinderwala, GC. New genome sequences were generated by TM, SS, SH, and DLS. RNA sequencing and genome annotation was performed by DLS. Computational analyses and figure preparation were performed by FBhinderwala and DLS. The project was conceived by AK, FBhinderwala, FBazan, AMG, and DLS. The project was funded by AMG and DLS. The manuscript was written by FBhinderwala, AK, AMG, and DLS. All authors contributed to manuscript revisions.

## Methods and Materials

### Tests for protein expression

Seven of the eighteen tested bicycle proteins could be expressed in sufficient amounts in either bacterial or Baculovirus-mediated insect cell expression systems for structural studies. Poor expression of the remaining bicycle proteins may have resulted from improper folding and/or degradation of these cysteine-rich proteins.

### Constructs

A synthetic gene sequence encoding the predicted secreted g2703 protein was codon optimized for expression in *E. coli* and purchased from IDT DNA Technologies. This DNA was inserted between NotI and BamHI restriction sites into the pET51b vector. (MilliporeSigma/Novagen) The final protein construct contains an N-terminal Strep-Tag II and a C-terminal His_10_ tag. The DNA encoding g3873 was purchased from GenScript and inserted into the pET28a vector (MilliporeSigma/Novagen) using Nde1 and Xho1 restriction sites. The resulting protein construct contains an N-terminal His_6_ tag followed by a TEV cleavage site.

### Protein Expression and Purification

For protein expression, *E. coli BL21(DE3)* was transformed with the pET51 vector encoding *g2703*. Cells were grown for 8 hours at 37°C in 5mL of LB medium containing 100 µg/ml carbenicillin. This starter culture was added to 50mL of modified M9 medium, containing 4g/L ^15^NH_4_Cl and 0.2 g/L glucose as nitrogen and carbon sources, respectively. Cells were grown overnight and diluted to an A_600_ of 0.25 into 1L of fresh modified M9 medium and grown to an A_600_ of 0.8. Expression was induced with 500 µM IPTG overnight at 18°C. For selenomethionine labeling, 100 mg/L of selenomethionine was added to the culture at an A_600_ of 0.5, cells were grown at 18°C for 1 hour, induced with 500 µM IPTG, and grown further overnight. Cells were harvested by centrifugation at 4000g at 4°C and resuspended in lysis buffer (50 mM NaH_2_PO_4_, pH 8, 250 mM NaCl, 1x EDTA-free protease inhibitor cocktail) and lysed on ice by sonication (for a total process time of 10 min with 5 sec on and 10 sec off at 50% power). Cell debris was removed by centrifugation at 16,000 g for 45 min at 4°C, and the Strep-tagged protein was purified over a StrepTrap HP column (Cytiva) in 50 mM NaH_2_PO_4_, pH 8, 250 mM NaCl, pH 8.0 (affinity buffer). The column was washed with 10 volumes of affinity buffer, and the bound protein was eluted in affinity buffer containing 250 mM desthiobiotin. Protein-containing fractions were pooled and concentrated using an Amicon-type centrifugal device with a 10kDa cutoff filter. 5mL of the concentrated sample was further purified by size exclusion chromatography (SEC) over a Sephadex 75 column (Cytiva) in SEC buffer (50 mM NaH_2_PO_4_, pH 7, 150mM NaCl). Following SEC, a final purification step was performed by ion exchange over a MonoQ column with a thirty-column volume gradient from 0-0.5M NaCl in 50 mM NaH_2_PO_4_, pH 7 buffer. Protein fractions at 0.15 M NaCl corresponding to the purified g2703 were pooled.

For protein expression of the His-tagged g3873 protein, Shuffle T7 cells (New England Biolabs) were transformed with the pET28a vector containing the g3873 gene. Cultures were grown as detailed above for g2703 expression. Harvested cells were resuspended in His-affinity buffer (50 mM NaH_2_PO_4_, pH 8, 250 mM NaCl, 30 mM imidazole) with 1x EDTA-free protease inhibitor cocktail and lysed by sonication as described above. Since most of the protein was in the insoluble fraction, the pellet was further sonicated in denaturing affinity buffer (50 mM NaH_2_PO_4_, pH 8, 250 mM NaCl, 30 mM imidazole, 8M Urea) and the supernatant was clarified by centrifugation. The cleared supernatant was loaded on two tandem 5 mL HisTrap FF columns (Cytiva), and the columns were washed with five column volumes of denaturing affinity buffer. The bound protein was refolded on the column by washing with 100 column volumes of a linear urea gradient from 8M urea to 0M urea in the affinity buffer. After a wash with five column volumes of 6% elution buffer (50 mM NaH_2_PO_4_, pH 8, 150 mM NaCl, 500 mM imidazole):94% affinity buffer, the refolded G3873 protein was eluted in 100% elution buffer. The N-terminal His tag was cleaved by adding 10:1 protein: TEV protease to the protein and incubation at room temperature for 4 hours. The cleaved g3873 protein was passed over a His-trap affinity column to remove the remaining TEV protease and the His-tag fragment. The column flow-through was further purified by size exclusion over a Sephadex 75 column in the SEC buffer. Protein-containing fractions were pooled and concentrated using an Amicon device with 10 kDa cut-off filters. Final protein concentrations were estimated by measuring A_280_ (NanoDrop Instruments), using an extinction coefficient of 15,600 M^-1^ cm^-1^ for g3873 and 49,800 M^-1^ cm^-1^ for g2703. Protein purity and labeling efficiencies were assessed by mass spectrometry. Pure protein samples were flash-frozen in liquid nitrogen and stored at -80 °C until further use.

### Crystallization and data collection

Both g2703 and g3873 proteins were buffer exchanged into 25 mM HEPES buffer at pH 7.0 and used for crystal screening at 4°C and 20 °C by the hanging drop procedure. Pure protein was mixed with an equal volume of reservoir solution. Crystallization hits obtained from initial screens were optimized using a range of pH, protein concentration, and varying buffer, salt, and precipitant composition. Diffraction-quality crystals were harvested, flash frozen in liquid nitrogen with cryoprotection in 20% (v/v) glycerol containing reservoir solution. Diffraction datasets were collected at the Stanford Synchrotron Radiation Lightsource (SSRL) BL12-2 beamline. The g2703 and the Se-Met g2703 crystals were in the space group of P2_1_2_1_2_1_ and diffracted to 1.4 Å resolution. The crystals of g3873 diffracted to 2.0 Å and were phased using sulfur-based anomalous diffraction. Crystallographic datasets were integrated and scaled using CCPN. Crystal data collection parameters and all structure statistics are summarized in Table S1.

### Genome sequencing

High molecular weight genomic DNA was prepared from samples of aphids collected from a single gall for each of the species *Hormaphis hamamelidis, Hormaphis betulae*, and *Hamamelistes spinosus* (Table S11) according to the manufacturer’s protocol (CG000145_SamplePrepDemonstratedProtocol -DNAExtractionSingleInsects.pdf, https://support.10xgenomics.com/genome-exome/sample-prep/doc/demonstrated-protocol-dna-extraction-from-single-insects). HMW DNA was quantified using a Qubit fluorometer (Thermo Fisher Scientific, Cat #Q32866) with the Qubit™ dsDNA BR Assay Kit (Thermo Fisher Scientific, Cat #Q32850) and fragment size was assessed by pulsed-field gel electrophoresis (PFGE). The DNA was run on a 1% agarose gel (Seakem Gold Agarose, Lonza, Rockland, ME, USA, Cat #50150) in 0.5x TBE buffer using the BioRad CHEF Mapper system (BioRad, Hercules, CA, USA, Cat #M1703650) for 15 hours. The gel was then stained with SYBR Gold dye (Thermo Fisher Scientific, Waltham, MA, USA, Cat #11494). PFGE results indicated that the aphid DNA ranged in size from 50 to 300 kbp

For all three species, 10× Genomics Chromium linked-read libraries were generated using the Chromium Genome Library Kit & Gel Bead Kit v2 (10× Genomics, San Francisco, CA, USA, Cat #120258), following the manufacturer’s protocol. The Illumina reads were assembled using the Supernova assembler (ver. 2.1.1).

For *Hamamelistes spinosus*, HiC reads were acquired to facilitate scaffolding of this genome. A single insect was frozen in liquid nitrogen in an Eppendorf tube, ground with a plastic pestle in 1 mL 1% formaldehyde for 20 min. Formaldehyde was quenched by adding 110 μL 1.25 M glycine for 15 min. The sample was spun at maximum speed for 15 min in a tabletop centrifuge. The liquid was removed and replaced with phosphate buffered saline, then spun again. The supernatant was removed, and the sample was stored at −80 °C prior to shipping to Phase Genomics (Seattle, Washington, USA) for Hi-C library preparation and sequencing.

We used a 3D-DNA assembly pipeline (65) in “haploid mode” to detect misjoins in the *H. spinosus* Supernova assembly and generate chromosome-scale super scaffolds. HiC alignments for the 3D-DNA pipeline were generated with Juicer v1.6.2 (66) using default settings. The scaffolded assembly was manually reviewed based on inspection of the HiC contact map using Juicebox Assembly Tools (67). The assembly was screened for contamination with BlobTools v1.0.1 (68, 69) using read coverage from the 10x Genomics linked reads and taxonomy information from BLASTN v2.2.31 (70) searches against the National Center for Biotechnology Information (NCBI) nucleotide database. We assessed the quality and completeness of the final assembly using BUSCO v3.0 (71, 72) with the arthropoda_odb9 gene set (73) and by comparing K-mer content of the 10X Genomics linked reads to the assembly with KAT comp v2.3.1 (74).

### RNA sequencing

For all three species, salivary glands were dissected from individual insects by placing insects in phosphate buffered saline and then grasping insects with forceps at the pronotum and anterior abdomen and pulling apart. The salivary glands remain attached to the brain on the anterior portion. The glands were isolated from other tissue using Minutien insect pins, with their tips bent ⍰30°, mounted in micro dissecting needle holders. Separated glands and the remaining carcass were collected separately into 100 μL of Arcturus PicoPure Extraction buffer. Total RNA was prepared using the Arcturus PicoPure RNA Isolation kit including the optional DNAse step. Barcoded RNASeq libraries were prepared for Illumina NextSeq 550 sequencing (150 bp PE) with a method described previously (75).

#### Annotation of bicycle genes

The new genome assemblies were re-annotated existing genome assemblies with RNAseq evidence using BRAKER (76). We then predicted *bicycle* genes using a gene-structure classifier (18) and manually annotated all predicted *bicycle* genes within Apollo (77) using RNAseq evidence collected from salivary glands. GFF files containing BRAKER annotation plus manual annotations are provided as supplementary material.

### Phylogeny estimation

An unrooted phylogeny was estimated for the seven species used in this study (Figure S4c) using *getphylo* (78) using all default parameters and starting with FASTA files of all predicted proteins from our new genome annotations for all species.

### AlphaFold2 protein structure prediction

AlphaFold2 (48) was installed on the Janelia Compute Cluster. Initial predictions for G3873 and G2703 were carried out using default sequence databases. Since these predictions did not match the crystal structures, we generated custom MSAs for bicycle proteins using predicted protein sequences for previously published aphid genomes and our new aphid genome sequences using a custom bash shell script (Supplementary Material). We then ran AlphaFold2 with the flag --use_precomputed_msas. These predictions, generated using custom MSAs, had higher confidence and resulted in models that were similar to the crystal structures for both proteins.

We used the above procedure to predict models of all potential bicycle proteins in *H. cornu, A. pisum*, and *S. chinensis*. In addition, we also predicted bicycle protein models for all proteins identified by *jackhmmer* search (79) with G2703.

### AF2 predictions and filtering

The initial set of all predicted models was filtered according to the following criteria: (1) low-quality AF2 predictions were removed, using a generous cut-off of median pLDDT score of 60 for at least 80% of the length of the protein sequence, and (2) the polypeptide had to contain at least 70 residues and more than one alpha helix (Supplementary Figure S11). From the initial set of 4093 proteins, applying the requirement of a median pLDDT >60 for at least 80% of the polypeptide length eliminated 1530 proteins, and the length and helical content filter eliminated another 140 models. The remaining 2423 protein models were inspected manually, and an additional 20 models were removed since these proteins exhibited >99.0% sequence identity to another bicycle protein; they were essentially duplicates. Details regarding the number of proteins per aphid species predicted by subsequent analyses are provided in the SI (Table S7)

### Sequence variability across the bicycle proteins

The variability in overall amino acid sequence space that is sampled by the bicycle proteins was examined using the length-normalized Levenshtein (50) distance. This provides a measure of the difference between two sequences. For each sequence pair, *a* and *b*, the Levenshtein distance *d* (*a, b*) was computed, which provides the minimum number of substitutions, insertions, and deletions required to transform *a* into *b*. Since the dataset includes proteins with up to six CYC motifs, it is essential to normalize this distance by the protein max length:

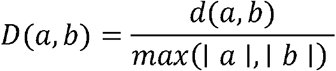

Where a and b denote lengths of the two protein sequences in the pair. This results in a symmetric distance between 0-1 for sequences of comparable length (with *D* = 0 for identical sequences) and permits us to compare divergence across the dataset. All pairwise distances were computed in an all-vs-all manner and used for the heatmap visualization. Identification of the most ‘isolated’ sequences within the bicycle proteins was carried out as follow: for each bicycle protein *a*, the fraction of other bicycle proteins *b* whose normalized distance from *a* exceeds a threshold *T* is calculated:

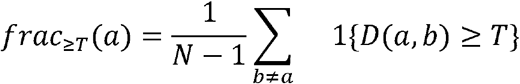

where *N* is the total number of bicycle proteins and 1 { *D* (*a, b*) ≥ *T* } is the indicator function. Large values of *frac*_≥*T*_ (*a*) indicate that *a* is distant from a large fraction of members in the dataset (i.e., an isolated sequence), whereas smaller values indicate that *a* is relatively close to most sequences.

We also generated an ABEGO Levenshtein distance map for all 2403 bicycle proteins in the filtered dataset. Comparing all 2403 bicycle proteins demonstrates that the ABEGO sequences within the family are highly diverse. We clustered the ABEGO sequences to illustrate sequence variation across the family and the lack of conserved sequence patterns within each aphid species. The dendrogram was colored at the lowest level using the species to which the corresponding bicycle protein belonged (Figure 3B)

### Evaluating physicochemical diversity across bicycle proteins

Physicochemical properties of all predicted bicycle models were computed based on the location and nature of each amino acid in the AF2 predicted structures. A total of 22 different features were evaluated (Table S8, 9). The solvent accessible surface area (SASA) was estimated using the FreeSASA algorithm (80), where all atom-level SASA were summed per residue to obtain the total SASA. Charged and hydrophobic SASA partitions were derived by defining hydrophobic (I, L, V, F, C, M, A, W, Y, P), positively charged (K, R, and H at pH 7.0), and negatively charged amino acids (D and E). f_SASA hydrophobic_ was calculated as SASA_hydrophobic_ / SASA_total_.

In addition, a set of features describing the charge density and net charge was calculated. The fraction of positive, negative, and hydrophobic residues was calculated by dividing individual values by the length of the sequence. Surface hydropathy, surface roughness, hydrophobic patchiness, amphipathic helical fraction, and max hydrophobic moment were also tabulated. All properties and the formulas used to compute them are described in detail in Supplementary methods.

The presence of correlations between these features was evaluated. Correlation will be present since some features contribute to the same overall property. We checked the range of each variable in order to prevent skewed assessment of the overall physicochemical nature of the proteins. The calculated parameters for all 2400 bicycle proteins are available in Table S9.

Using the reduced set of uncorrelated features, our analysis used a custom Python script within the *sklearn* toolkit (67) and the data were plotted using *matplotlib* (68). Non-linear dimensionality reduction was performed with t-SNE (51) and UMAP (52, 69) for all non-protein size-dependent parameters, i.e., 11 out of the 22 original and a reduced set of physicochemical parameters (9 out of 11), with a correlation threshold set at 0.85. After assessing various combinations of parameters, we opted to visualize the UMAP projection using 30 neighbors and a distance of 0.05, with a Leiden resolution of clusters at 1.2. This allowed us to preserve the global structure of the map and maximize the consistency of cluster memberships. Medoids representing each of the clusters were identified and used to highlight the physicochemical properties of members in each cluster. The overall distribution of each feature across aphid species and Leiden clusters is provided in the supplementary data (Figure S16).

### Mapping the diversity of structural space across bicycle proteins

Using the 2400 AF2 predicted bicycle protein models, we performed an all-versus-all comparison of TM scores (70, 71). This all-versus-all tm-score matrix was reduced to a two-dimensional space using t-SNE (51), with subsequent Leiden clustering (49). This resulted in a global map of predicted bicycle protein structures (provided in Figure 3B). Medoids representative of each cluster were plotted (Figure 4). Clustering was based solely on structural features, since no cluster over-represented a particular aphid species or a defined polypeptide chain length.

