## SupplementaryInformation for "Rapidly evolving aphid gall effector proteins exhibit saposin-like folds"

### Supplementary information for Rapidly evolving aphid gall effector proteins exhibit saposin-like folds

Fatema Bhinderwala: 0000-0002-3033-8438

Aishwarya Korgaonkar: 0009-0004-4244-0140

Kota Gopalakrishna: 0000-0001-9275-7059

Thomas Mathers : 0000-0002-8637-3515

Shuji Shigenobu: 0000-0003-4640-2323

J. Fernando Bazan : 0000-0002-3645-9935

Saskia Hogenhout : 0000-0003-1371-5606

Guillermo Calero: 0000-0003-3730-4676

Angela Gronenborn: 0000-0001-9072-3525

David L. Stern: 0000-0002-1847-6483

**e-mail**

Fatema Bhinderwala:

Aishwarya Korgaonkar:

Kota Gopalakrishna:

Thomas Mathers:

Shuji Shigenobu:

J. Fernando Bazan:

Saskia Hogenhout:

Guillermo Calero:

Angela Gronenborn:

David L. Stern:

### Table of Contents:

Supplementary Methods

**List of Supplementary Figures:**

**Figure S1** Mass spectrometry data of recombinant proteins.

**Figure S2:** Poisson-Boltzmann electrostatics surface of g3873.

**Figure S3:** Sulfur density in the anomalous diffraction map of g3873.

**Figure S4:** G2703 paralog family

**Figure S5:** Volcano plot of differential expression analysis of genes enriched in salivary glands versus body of gall foundresses of *H. cornu.*

**Figure S6:** G2703 MSA

**Figure S7:** Structural similarity between the N-terminal and C-terminal half of g3873 and saposin-like proteins in the PDB.

**Figure S8:** Structure comparison between CYC domains of G3873 and G2703 to the most similar saposins.

**Figure S9:** Structural similarity between the N-terminal and C-terminal half of g2703 and saposin-like proteins in the PDB.

**Figure S10:** AF3 and ESMFold predictions for g3873 and g2703 using single amino acid sequences.

**Figure S11:** AF2 prediction accuracy for all bicycle proteins across seven different aphid species using our custom MSAs.

**Figure S12:** AlphaFold2 prediction with custom MSA for G2703 and G3873.

**Figure S13:** Effect of downsampling MSAs on AF2 prediction accuracy.

**Figure S14:** Species distribution within each Leiden cluster of the t-SNE from Figure 4B.

**Figure S15:** Bar graphs depicting the proportional contribution of proteins from each species to each Leiden cluster in the t-SNE plot from Figure 4B.

**Figure S16:** Physicochemical signatures in space filling representations of Leiden cluster medoid models

**Figure S17:** UMAP of physicochemical signatures for bicycle proteins of different aphid species

**Figure S18:** Bar graphs depicting the proportional contribution of proteins from each species to each Leiden cluster in the UMAP plot from Figure 5.

**Figure S19:** Violin plots of physicochemical properties by cluster and species.

**Figure S20:** Shannon entropy of bicycle protein diversity mapped onto a ribbon representation of g3873.

**Figure S21:** Summary of disulfide topology for bicycle proteins and other disulfide containing proteins.

**Figure S22:** Frustration analysis of *H.cornu* bicycle proteins.

**List of Supplementary Tables:**

**Table S1**: Expression and crystallization results for bicycle genes tested for recombinant protein expression

**Table S2**: Crystal data collection parameters and structure statistics for g2703 and g3873.

**Table S3:** Disulfide bond conformations in the X-ray structure of G3873.

**Table S4**: Foldseek hits for g3873 and g2703 identify only poor quality matches (.xlsx file).

**Table S5**: Cα RMSD values between all known saposin structures in the PDB and the saposin-like domains in G3873

**Table S6:** TM scores for all known saposin-like proteins in the PDB and the saposin-like domains in G2703

**Table S7:** Statistics of AF2-predicted models for all seven aphid species

**Table S8:** List of physicochemical features

**Table S9:** Physicochemical properties for 2400 Bicycle proteins

**Table S10:** Metrics of structural space

**Table S11:** Genome sequencing statistics

**Supplementary Methods:**

**Materials:** All general chemicals were purchased from Fisher Scientific. Selenomethionine for labeling was purchased from Millipore Sigma and all crystallography screens and materials to prepare custom crystallization solutions were purchased from Hampton Research.

**Mass spectrometry for recombinantly expressed proteins:**

All ESI LC-MS measurements were performed at a protein concentration of 1 μM on a Bruker Q-TOF instrument, using a reverse-phase AdvanceBio peptide guard column (Agilent Technologies), with mobile phases A and B comprising 5% acetonitrile with 0.01% FA and 80% acetonitrile with 0.01% FA, respectively. The resulting LC-MS spectra were processed using Bruker Compass Software, and the MS data were processed using Maximum Entropy-based deconvolution to obtain the M+ ion mass for each sample. The instrument was calibrated using the ESI Low Tuning mix I (Agilent Technology) to a 1.0 ppm mass % difference before each use.

**Ellman's assay to determine free sulfhydryl groups in G3873**

Purified protein was buffer exchanged into 0.1 M sodium phosphate buffer, pH 8, 1 mM EDTA and concentrated to 120 μM prior to use in the Ellman's assay. A working solution of Ellman's reagent (DTNB) at 10 mM concentration in 0.1M sodium phosphate buffer, pH 8, was freshly prepared. A six-point calibration curve was prepared using reduced L-glutathione and L-cysteine over a linear range from 10-200 μM of total thiol. Assays were performed in a 96-well plate format by mixing 5 μL of sample with 95 μL of DTNB working solution (three technical replicates per condition). Plates were incubated at room temperature for 10 min, and absorbance at 412 nm was measured using a Tecan Spark spectrophotometer. Total free sulfhydryls were calculated using Beer-Lambert's law using the molar extinction coefficient of DTNB of 14,150 $M^{-1}$ $cm^{-1}$.

**Calculation of disulfide bond conformation and bond energies in G3873**

Disulfide-bond geometry was analyzed with the UCLA Disulfide Bond Dihedral Angle Energy Server by submitting the PDB coordinates of the experimental structure of G3873. For each disulfide linkage, the server then calculated the five defining cystine dihedral angles, $\chi1$, $\chi1'$, $\chi2$, $\chi2'$, and $\chi3$, and the corresponding empirical dihedral energy (1). Dihedral energy was calculated in kJ/mol using the following equation:

$$E = 8.37(1 + \cos 3\chi1) + 8.37(1 + \cos 3\chi1') + 4.18(1 + \cos 3\chi2)$$
$$+ 4.18(1 + \cos 3\chi2') + 14.64(1 + \cos 2\chi3) + 2.51(1 + \cos 3\chi3)$$

The resulting dihedral angles and energies for each disulfide bond are reported in Table S3 and were used to assess relative torsional strain and conformational favorability within the structure.

**AlphaFold3 and ESMFold predictions for bicycle proteins:**

Protein models were generated using the respective web servers for AF3 (2) and ESMFold (3). For each protein of interest, the corresponding amino acid sequence was submitted to the server using default settings. Predicted models were downloaded, and the respective per-residue confidence scores (pLDDT) were retained as B-factor. The top-ranked model (as returned by the server) was used for comparisons throughout.

**DALI and FoldSeek search for similar proteins using experimental structures for G3873 and G2703:**

DALI protein structure comparisons were performed using the web server (http://ekhidna.biocenter.helsinki.fi/dali) (4) with the crystal structures of both G3873 and G2703 as input models. Each structure was queried against the entire PDB to match at minimum 30 residues in length. Neither bicycle protein resulted in hits with a DALI Z-score >8, which would indicate similar fold or structural topology. A Foldseek search for both crystal structures of G3873 and G2703 as input PDBs was also performed using the default parameters on the web server (https://search.foldseek.com/search) (5). These parameters are defined as 3Di+AA local alignment (alignment mode 2), sensitivity -s 9.5, E-value threshold 0.001, max 1000 hits, database pdb100 (redundancy-filtered PDB at 100% sequence identity / ≥95% coverage), iterative search off, cluster search off (representatives only). The results of the Foldseek search are summarized in supplementary Table S4.

**Calculation of backbone RMSD values and TM-score between the X-ray structures of G3873 and G2703 and all known saposin fold proteins in the PDB:**

Structural similarity between the N- and C-terminal CYC domains of G3873 and G2703 and all 52 PDB deposits with at least one saposin-like domain (classified by InterPro)(6) was assessed for individual chains. Each structure was compared to G3873 and G2703 by running TM-align for every pair. The TM-scores reported by TM-align (7) between the two structures were normalized to the length of the target protein. For each deposit-target pair, backbone RMSD was also computed.

**Shannon's entropy and sequence conservation**

Sequence diversity was quantified using Shannon entropy computed from a MSA. For each alignment column, the amino acid frequency was calculated across all sequences and converted into a per-position Shannon entropy score. Protein sequences were first filtered by length, and sequences ranging from 120–300 were retained. An MSA was generated from the filtered sequences using MAFFT. Gaps were excluded from frequency calculations, and positions with insufficient coverage were omitted from the summary statistics.

**Shannon entropy per MSA column (bits), computed over 20 amino acids only (gaps ignored).**

Formula:

$$H_j = -\sum_{a \in AA20} p_j(a) log_2 p_j(a)$$

Where: $p_j(a) = n_j(a)/N_j$, with $N_j$ = number of non-gap AA 20 characters in column $j$.

The first sequence in the alignment was used as the reference to map column entropies onto residue positions. To visualize entropy on the structures, the reference sequence was globally aligned to G3873, and entropy values were written into the B-factor field of matched residues.

**Heatmaps and dendrograms for ABEGO Levenshtein distances for bicycle proteins**

*Sequence collection and preprocessing*: Protein sequences were compiled into a FASTA file (n=2403) with identifiers encoding protein name and length (e.g., hc_g3873|chain=A|len=169). All sequences with >0.99 sequence identity were removed from downstream analyses.

*ABEGO string generation*: ABEGO is a 5-letter code for protein backbone conformation that bins backbone dihedral angles ϕ and ψ for each residue into a small number of common regions in the Ramachandran map (8): A corresponds to the right-handed α helical basin, B and E correspond to the extended/β strand-like basin, G corresponds to the left-handed α helical basin favored by glycine residues, and O captures outlier conformations. We used the following implementation: G (αL/left-handed region; $0 \leq \phi \leq 120$ and $-60 \leq \psi \leq 90$), A (αR/right-handed region; $-120 \leq \phi \leq -20$ and $-90 \leq \psi \leq 50$); B (β region; $-180 \leq \phi \leq -90$ and $\psi \geq 90$ or $\psi \leq -120$), E (extended region; $-180 \leq \phi \leq -60$ and $50 \leq \psi \leq 180$), and O (other/undefined: all remaining (ψ) combinations.

*Visualization:* For each distance matrix, heatmaps were generated using a continuous color map with low values indicating greater similarity and high values indicate greater dissimilarity. The lower bound of the color maps was changed to enhance contrast among intermediate distances, depending on whether the raw or ABEGO distances are displayed.

Hierarchical clustering was performed using average-linked agglomeration applied to pairwise distances, and the lowest tiers of the dendrogram were colored to correspond to the species in which the genes were found.

**Calculating and mapping physicochemical properties**

A total of 22 physicochemical features were computed for each protein model (Table S7). These properties combine both sequence-derived and structure-derived properties that capture composition, hydrophobicity, electrostatics, and amphipathic character. Solvent-accessible surface area (SASA) was computed using FreeSASA (9).

Atom-level SASA values were combined to yield residue-level SASA and summed across all residues to obtain the total SASA. Residues were assigned to three classes: hydrophobic, positively charged, and negatively charged, and SASA values were summed within each class to obtain SASA_hydrophobic, SASA_pos and SASA_neg. The hydrophobic surface fraction (a potentially relevant property that can regulate saposin-like activity and protein-protein interaction) was reported as a frac_sasa_hydrophobic. In each case, the residue was considered surface-exposed for these calculations if the SASA was >5Å.

*Surface hydropathy*: Surface hydropathy was quantified using the Kyte-Doolittle scale, and two properties were calculated: a SASA-weighted mean KD value and a median KD value over the exposed residues.

*Surface roughness*: A global proxy of surface roughness was computed as the ratio between total SASA to the surface area of the 3D convex hull using the SciPy (10) ConvexHull function.

*Surface hydrophobic patchiness*: Hydrophobic patches were identified by clustering the exposed hydrophobic residues using sidechain centroid coordinates and the Density-based special clustering of applications of noise (DBSCAN) (11). This yields three parameters: the number of hydropatches, the mean size of the patch, and the maximum size of the patch.

*Charge and charge-density*: Sequence level composition was computed as the fraction of positively charged, negatively charged, and hydrophobic (FLIV) residues. This was combined with net charge, exposed net charge, and exposed net charge over SASA.

*Surface charge dipole*: To capture spatial polarization of surface charge, a SASA-weighted dipole vector was computed from exposed charge residues.

*Amphipathic helical content*: The maximum hydrophobic moment was computed as the largest hydrophobic moment value observed over a helix, and an amphipathic helical fraction was calculated as a hydrophobic moment > 0.35 over a helix.

**TM-scores between AF2-predicted models of bicycle proteins from Hormaphis cornu and the X-ray structures of G3873 and G2703**

The TM-scores between all reliably predicted (median pLDDT >60 over 80% of the residues) bicycle proteins in *Hormaphis cornu* (n=243, for sequence lengths of 120-300 residues) were computed using the experimental X-ray structures of G3873 and G2703 as targets. The resulting TM-scores were visualized as scatter plots to illustrate the structural diversity in the set.

**All-vs-all TM-score matrix for all reliable AF2-predicted models of bicycle proteins of seven aphid species**

An all-vs-all TM-score matrix using the TM-align tool was calculated from 2400 models, running TM-align for each pair (i, j). Two TM-scores are obtained for each comparison, one normalized by the length of each of the two polypeptide chains (i and j). The TM-scores were converted into a distance matrix, with structurally similar proteins exhibiting small distances (high TM-scores). A kNN graph constructed from the TM-score-derived distances connects each protein to its nearest neighbor. Community grouping was performed using the weighted KNN graph using the Leiden algorithm (12). The resulting structural similarity space was embedded into two dimensions using t-SNE (13)

**Medoid selection in physicochemical and the structural space**

For each Leiden cluster $C$, a representative medoid structure was selected as the member with minimal mean distance to all other members of the same cluster:

$$m = argmin_{i \in C} \frac{1}{|C|-1} \sum_{j \in C, j \neq i} D_{ij}$$

The medoid maximizes the average TM-score compared to other cluster members. Structural models of medoids were used as representatives for cluster-specific visualization.

**APBS electrostatic surfaces**

Electrostatic surfaces were generated for AF2-predicted medoid models. For each medoid coordinate set, atomic charges and radii were assigned using pdb2pqr. Protonation states were assigned at pH 7.0 using the AMBER forcefield and outputs written as .pqr files for

APSB input. Electrostatic potentials were computed using APBS (14, 15) using the linearized Poisson-Boltzmann equation, a protein dielectric of 2.0 and a solvent dielectric of 78.0. The solvent probe radius was set to 1.4 Å, the surface density was set to 10.0, and the ionic concentration was set to 0.15 M of monovalent ions. Electrostatic maps were written as .dx grids and used to generate structure frames using a fixed color ramp of −5 to +5 kT/e in PyMOL (16).

**Disulfide topology across aphid bicycle proteins:**

HIgh confidence bicycle protein models were screened for disulfide bonds by identifying all pairs of cysteine sulfur atoms (Sγ–Sγ) within 3.0 Å of one another, where the threshold allows us to capture both predicted disulfide bonds and near-miss geometries of two cysteines occupying proximal positions in the predicted fold. Within each model, cysteines were renumbered 1→N in N-terminal to C-terminal order; the resulting set of bonded cysteine pairs was recorded as a sequence of index pairs and any cysteine without a partner within the cutoff was retained in the count as a free cysteine. Each model was then assigned to a topology category based on the geometric arrangement of its bonded skeleton: concentric-nested (e.g., 1-4, 2-3 where one disulfide bond is enclosed by another with no crossovers), spanning + adjacent (e.g., 1-6, 2-3, 4-5, where one long-range bond plus adjacent inner pairs exists), all-adjacent (e.g., 1-2, 3-4 where all sequential cysteine are bonded), and incomplete (any model with one or more cysteines unaccounted for at the 3.0 Å threshold). Free cysteines in incomplete models were further sub-classified as would-be-bonded, near-miss, scaffold-proximal, or orphan based on the distance from each unpaired cysteine Sγ to its nearest cysteine Sγ partner to avoid missed disulfide bonds due to poor side-chain geometry in predicted models.

**Frustration analysis and surface repatterning of frustrated patches:**

Two complimentary frustration analyses were performed on the three dominant Cys disulfide topology subgroups (n=133) identified from the disulfide topology analysis, restricted to the *H.cornu* Bicycle proteins. For each subgroup, the cleaned Foldmason structural MSA (17) and matched AlphaFold-predicted single-chain models were submitted to the FrustraEvo web server (18) which computes per-structure local frustration (19) and aggregates the results per MSA column. Residues with single-residue frustration index $\leq -1$ were classified as minimally frustrated, $\geq +0.78$ as highly frustrated, and otherwise neutral. To test whether highly frustrated residues are spatially enriched at the protein surface (20), per-residue solvent accessible surface area (SASA) was computed for each model using the Shrake-Rupley algorithm (21) as implemented in FreeSASA (9) with Bondi van-der-Waals radii and a 1.4 Å probe. Per-residue frustration index and SASA were merged on residue number within each model and pooled across all 133 models (25,730 residue observations). Spearman rank correlation of frustration index versus SASA was computed per subgroup and on the pooled dataset; residues were binned into per-subgroup SASA deciles, and enrichment of highly frustrated residues in the buried (lowest) versus solvent-exposed (highest) decile was tested by Fisher's exact test.

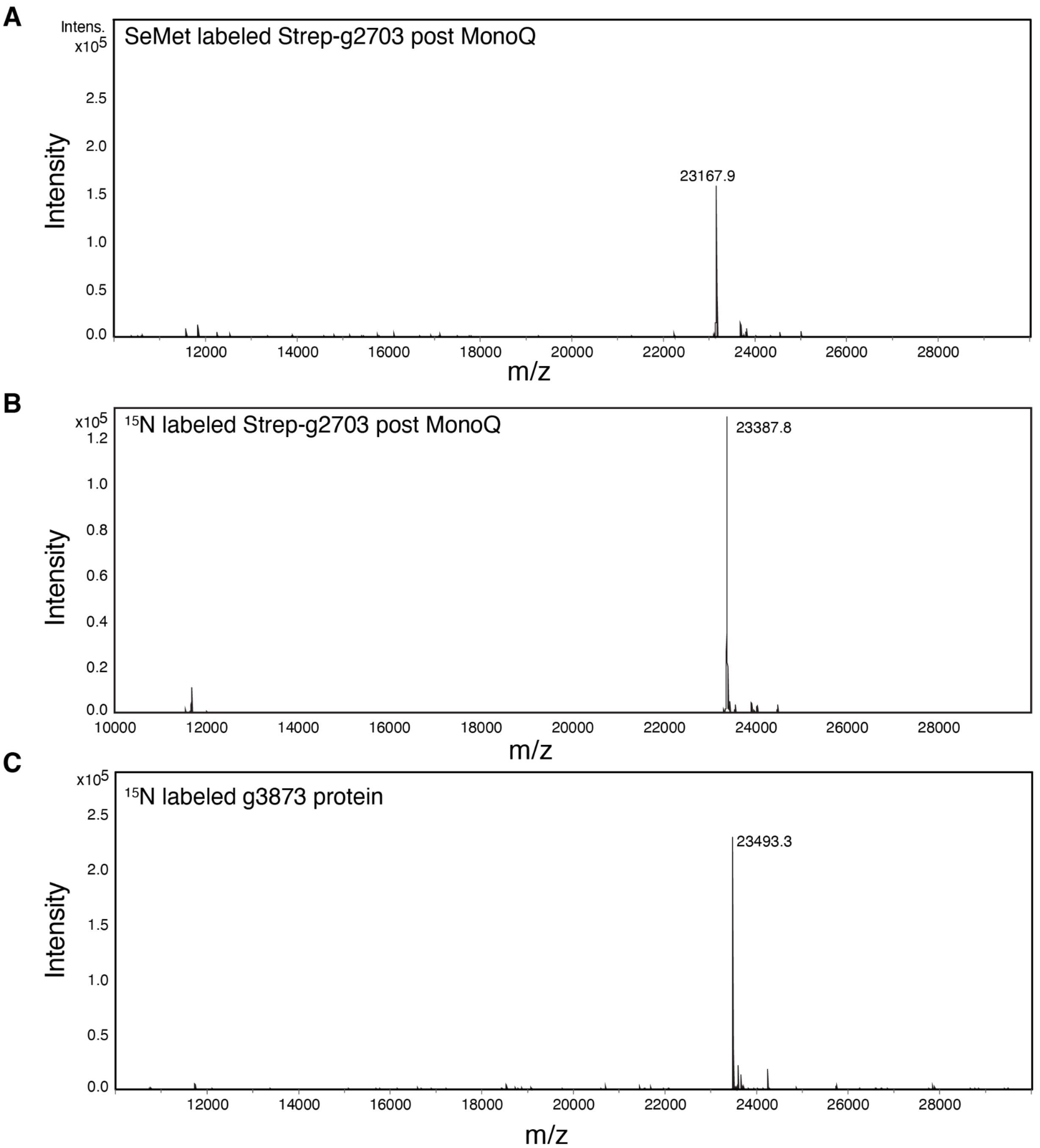

**Figure S1**: (A and B) ESI Mass spectrometry data of selenomethionine (SeMet) labeled Strep-tagged G2703 and 15N-labeled strep-tagged G2703 after cleavage of the tag. The calculated predicted masses for the full-length protein without the C-terminal 10x His-tag is 23387 Da. The predicted calculated mass is 23169 Da for the protein with a single SeMet residue.
(C) ESI Mass spectrometry data of cleaved $^{15}$N-labeled G3873 protein. The predicted calculated mass is 23490 Da.

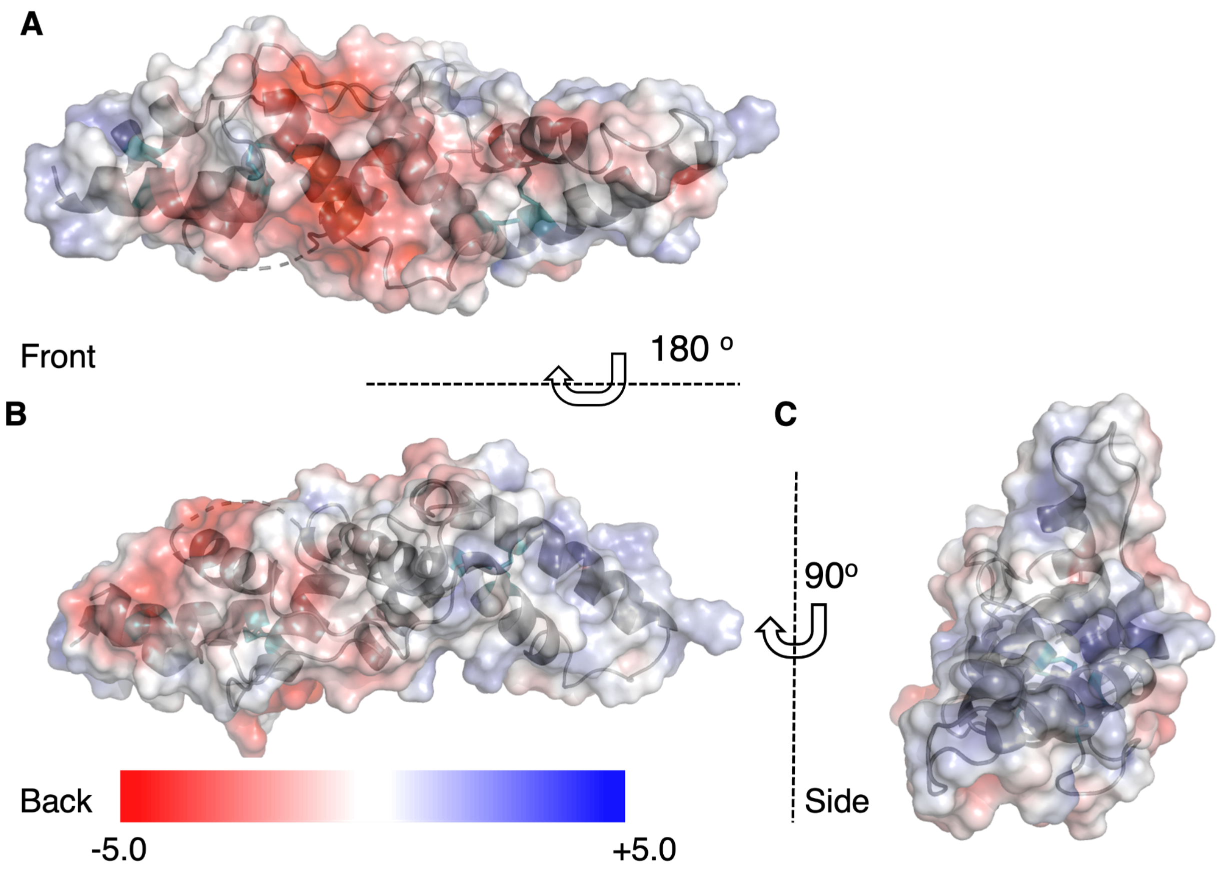

**Figure S2:** (A) Front, (B) back, and (C) side view of G3873 in space filling representation (gray) with the Poisson-Boltzmann electrostatic surface colored from red to blue for -5.0 to +5.0.

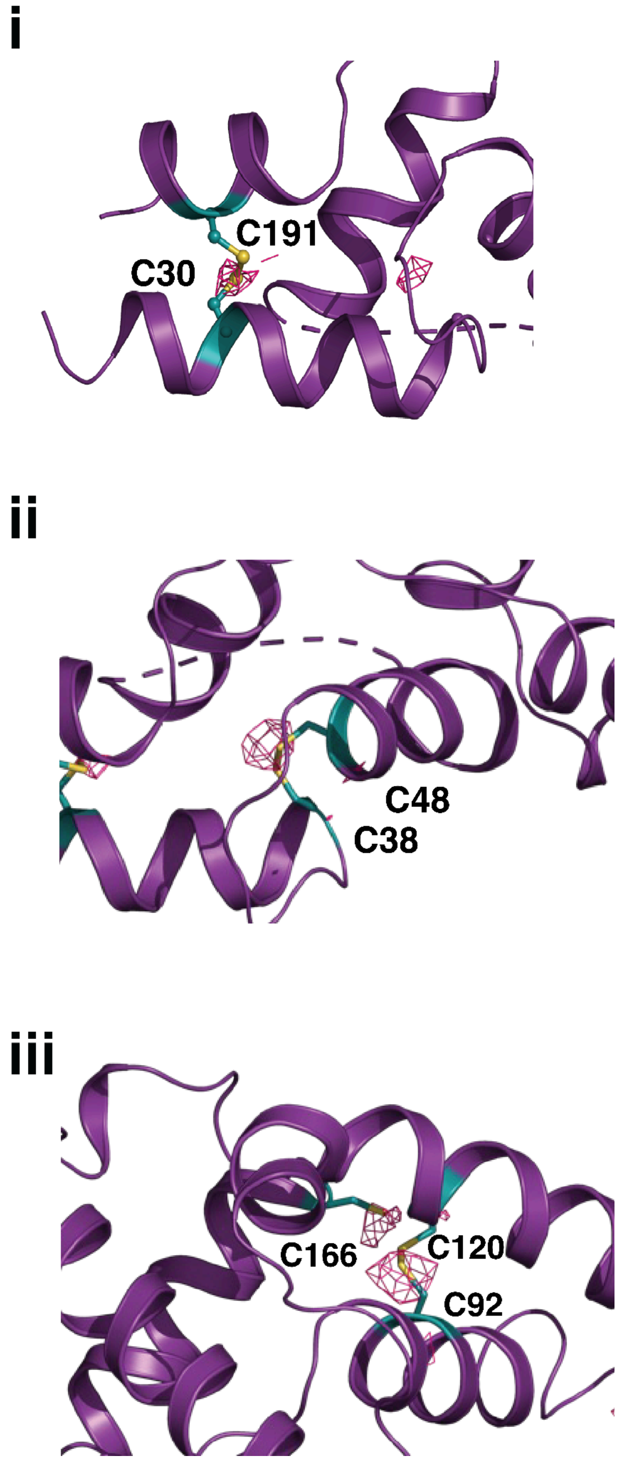

**Figure S3:** Sulfur density in anomalous diffraction map (magenta) at each of the three disulfide bonds (cyan and yellow ball and stick representation) in G3873 (purple ribbon representation) for (i) C30-C191, (ii) C38-C48, and (iii) C92-C120.

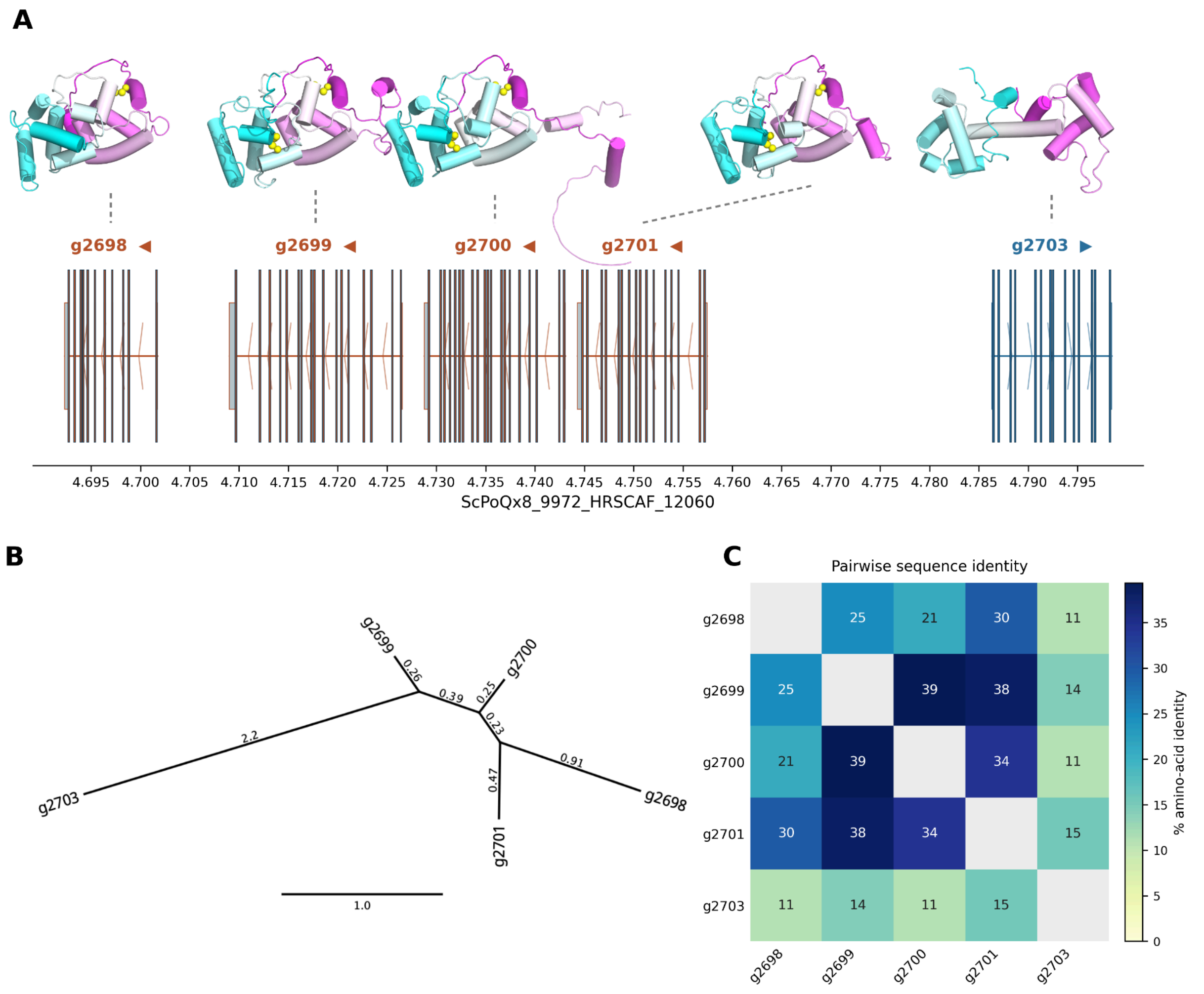

**Figure** S4:. Putative *bicycle* gene paralog family that includes *g2703* shows extreme sequence diversity.

(A) Gene models for five putative *bicycle* gene paralogs of *g2703* are found within an approximately 100kb interval. Vertical lines represent exons and the arrows indicate the direction of transcription. AF2 predicted protein structural models are shown above each gene model. All of these predicted protein structures include tandem saposin-like folds, with N-terminal domain (cyan) shown to the left of C-terminal domain (magenta).

(B) G2703 is the most divergent paralog of the family, with an estimated more than two substitutions per residue differentiating it from these paralogs.

(C) Pairwise sequence identity amongst these putative paralogs reveals little sequence similarity between G2703 and the other proteins.

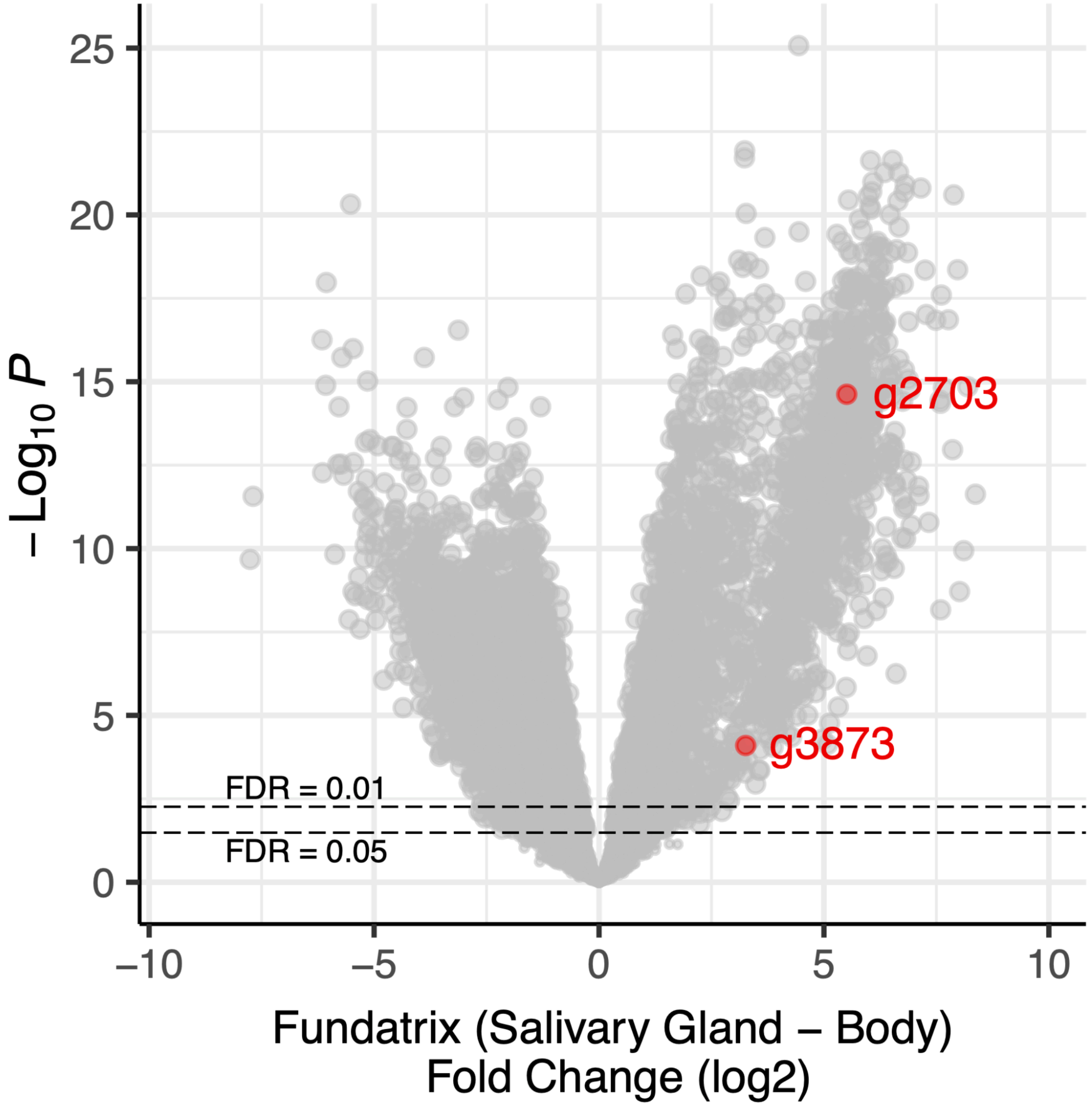

**Figure S5:** Volcano plot of differential expression analysis of genes enriched in salivary glands versus body of gall foundresses of *H. cornu.* The genes *g2703* and *g3873* are highlighted. The data and analysis pipeline are found in publication (22).

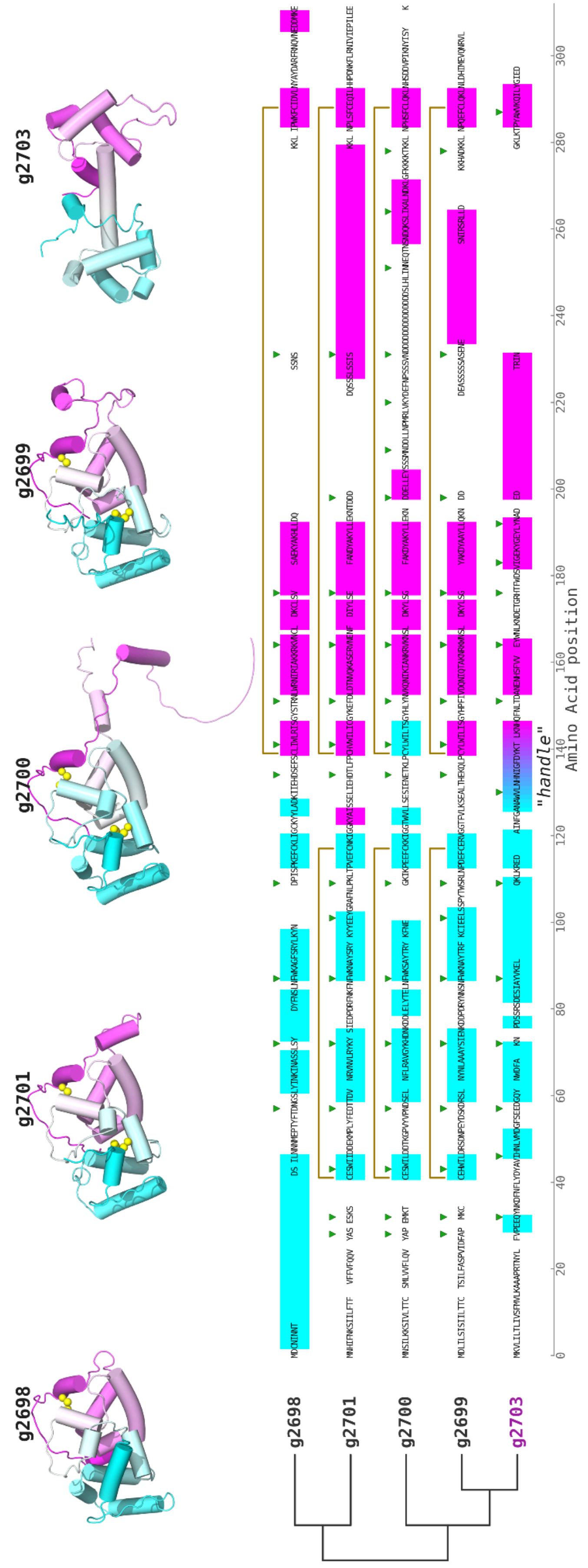
g2698
g2701
g2700
g2699
g2703
"handle"
Amino Acid position
0
20
40
60
80
100
120
140
160
180
200
220
240
260
280
300

**Figure S6**: Exon-aware multiple sequence alignment (15) of putative G2703 paralogs suggests that the cysteineless G2703 evolved from ancestors that contain disulfide bonds that link cysteines within the same saposin-like domain. In addition, the alpha-helical “handle” that connects the two saposin-like domains of G2703 likely evolved from a fusion of the last and first alpha helices of the first and second saposin-like domains, respectively. The AF2 predicted protein structures are shown at the top with saposin alpha helices indicated by cyan or magenta and disulfides indicated with yellow balls. The respective alpha helices shown on the MSA in colored blocks. Green triangles indicate locations of introns, and reveal extensive diversity in exon presence or absence amongst these closely related genes. Disulfide linkages between cysteines are indicated by gold colored brackets. The “handle” helix of G2703 is indicated.

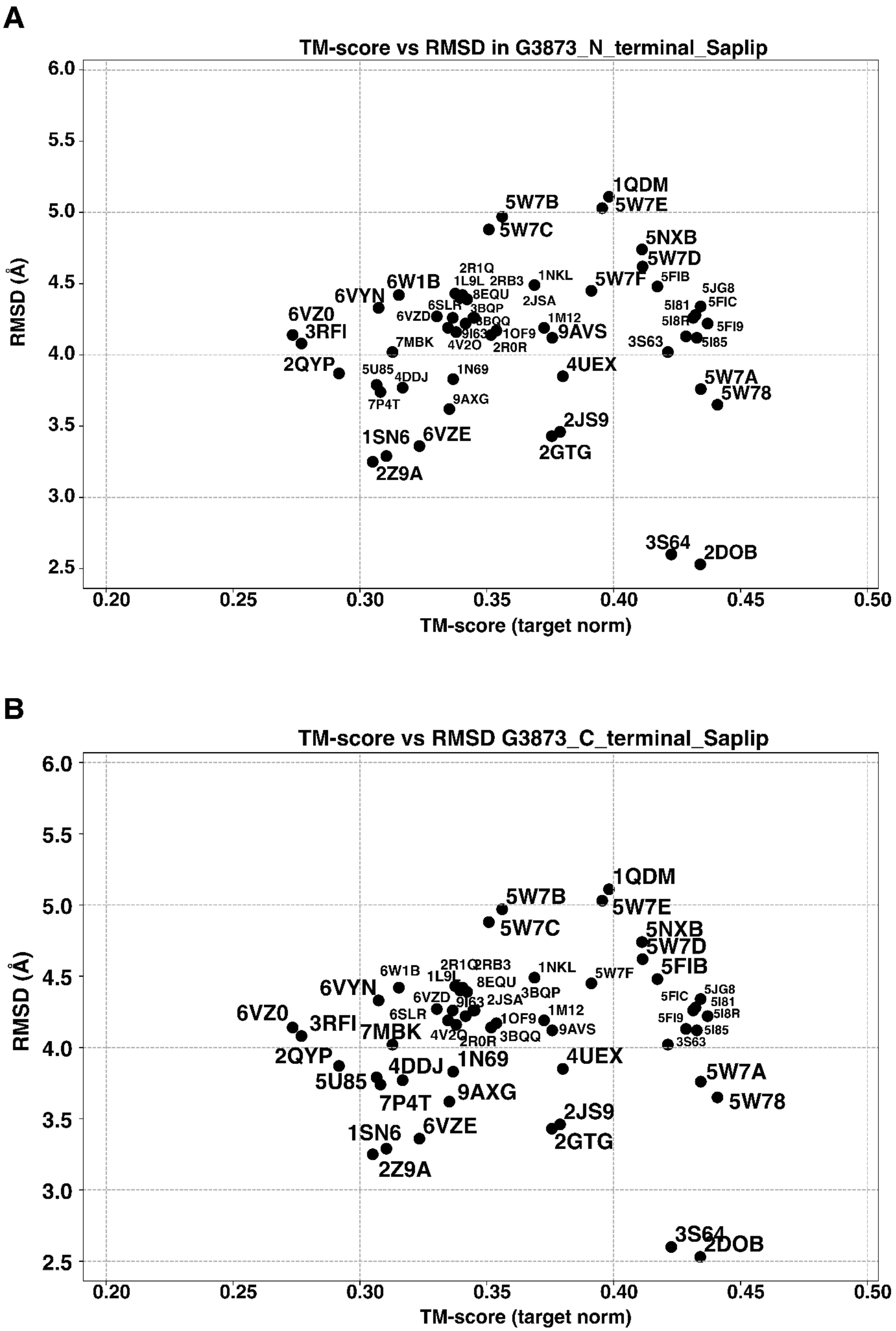

**Figure S7** Plots of RMSD values for the saposin N-terminal (A) and C-terminal (B) CYC motif domains of G3873 versus saposin-like proteins in the PDB.

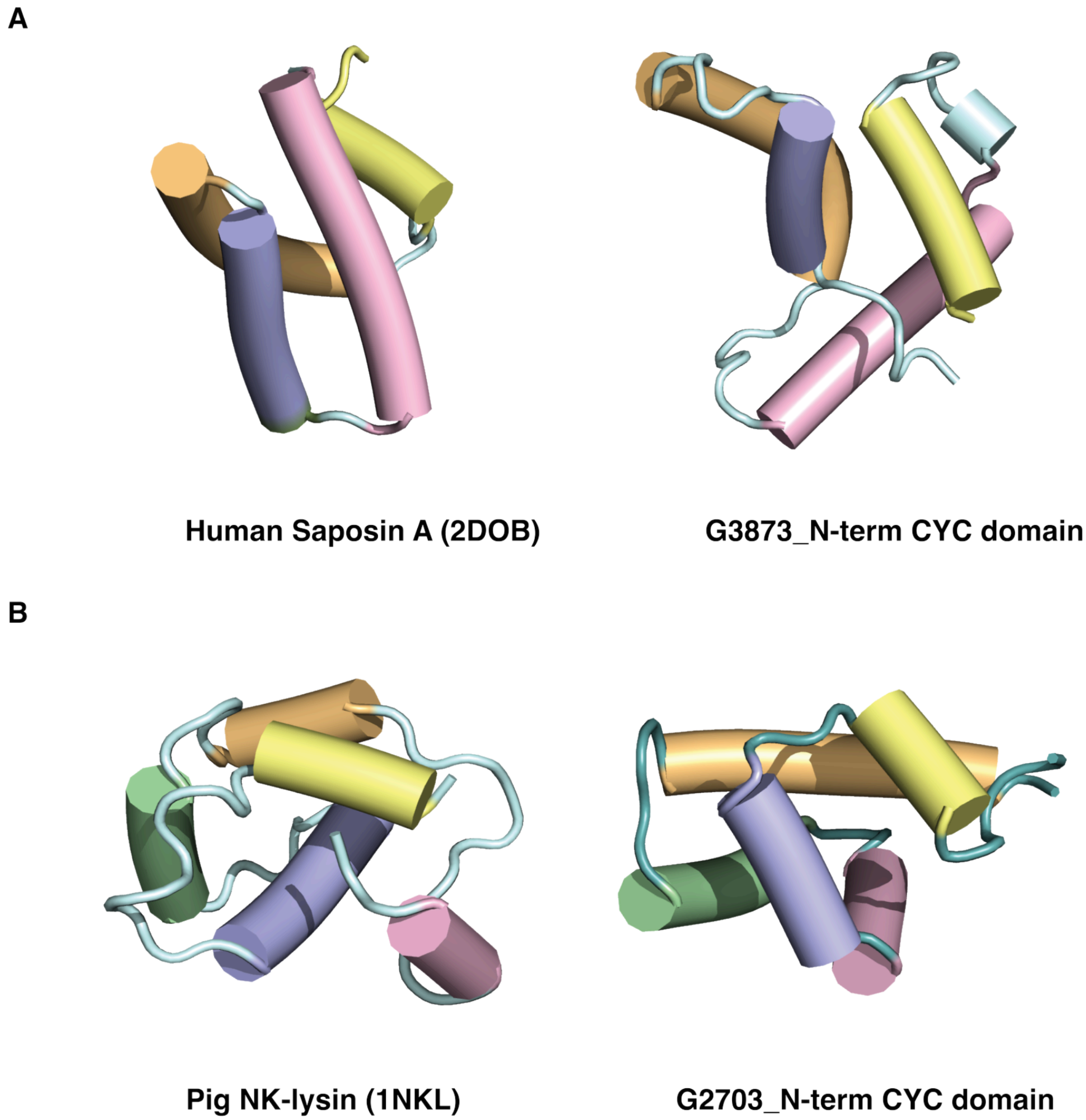

**Figure S8:** Structure comparison between the four-helix bundles of saposin-like proteins and bicycles CYC domains.

(A) N-terminal domain of g3873 next to human saposin A (PDB ID: 2DOB).

(B) N-terminal domain of g2703 next to pig NK-lysin (PDB ID: 1NKL). Helices are represented as cylinders and colored to show structural similarities

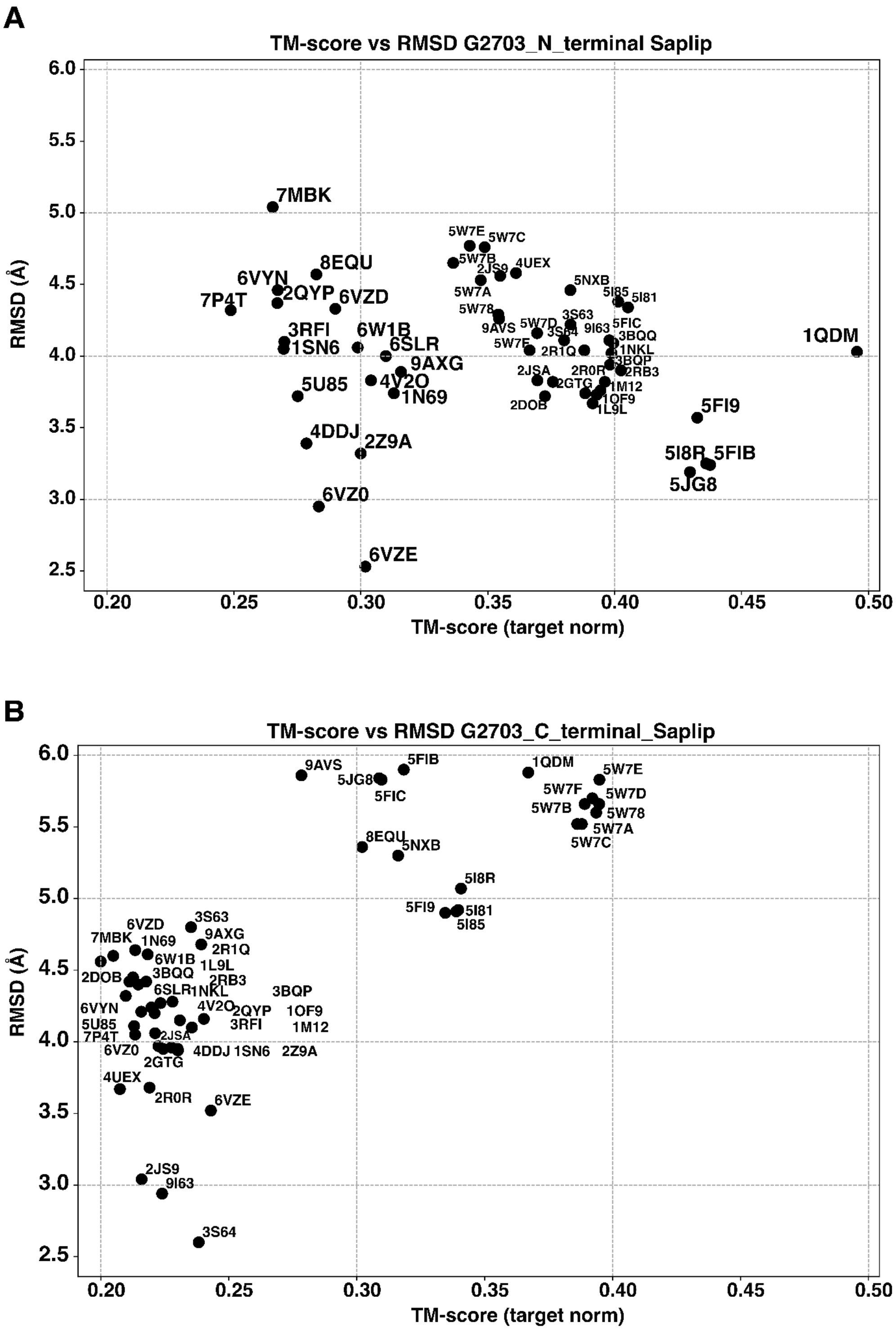

**Figure S9:** Plots of RMSD values for the saposin N-terminal (A) and C-terminal (B) CYC motif domains of G2703 versus saposin-like proteins in the PDB.

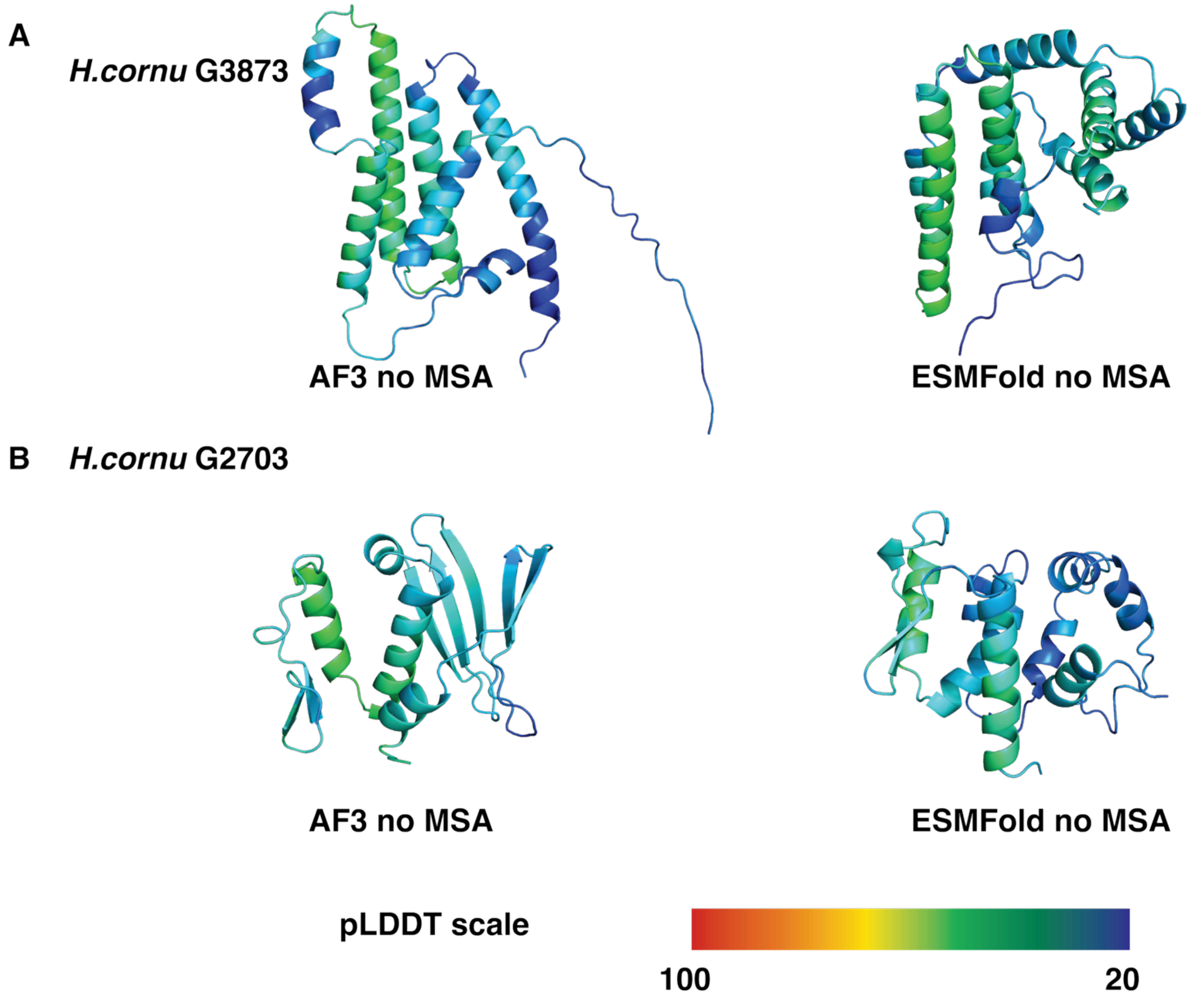

**Figure S10:** AF3 and ESMFold predicted models for G3873 (A) and G2703 (B) using a single amino acid sequence. The ribbon diagrams are colored according to their pLDDT scores (bottom bar).

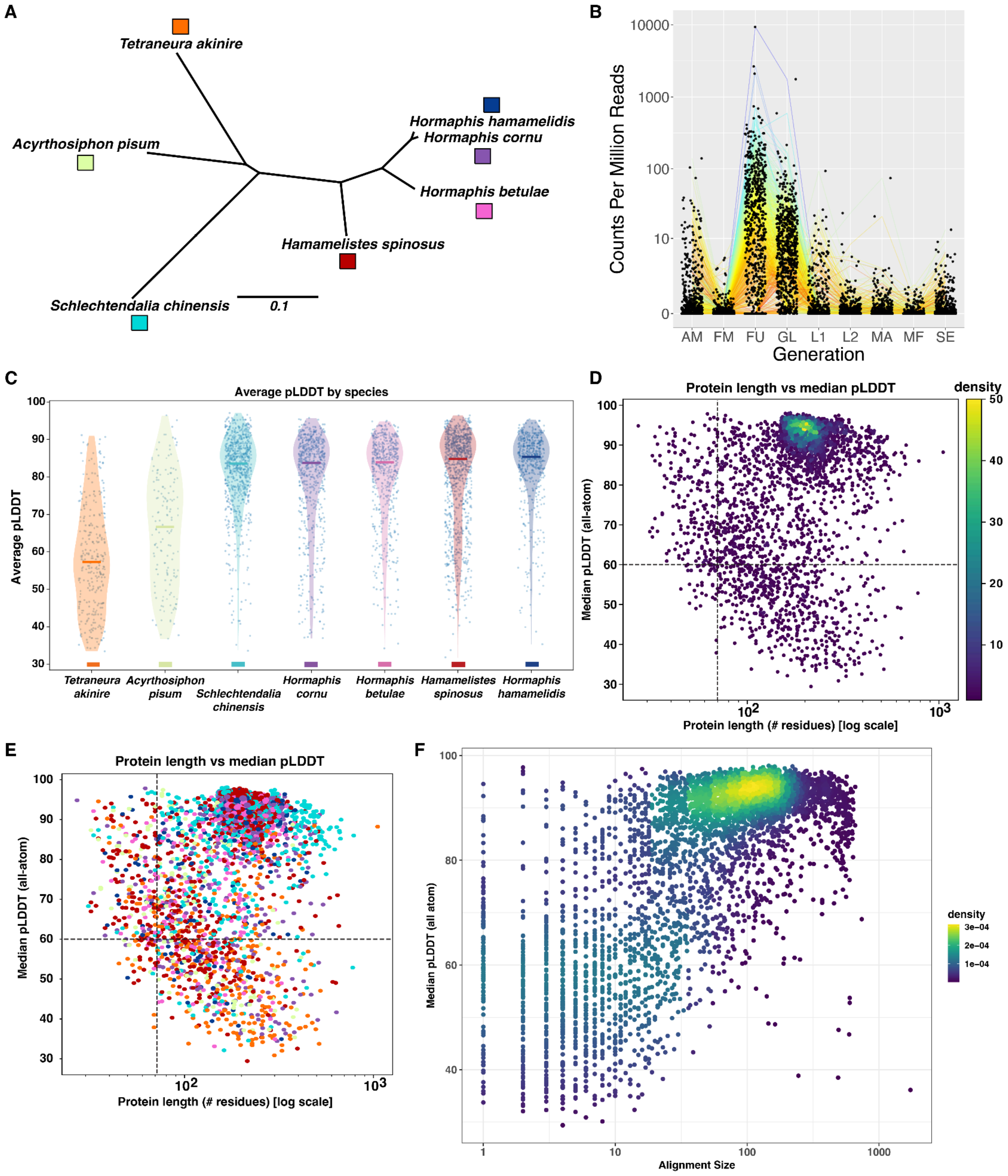

**Figure S11:** Distribution of pLDDT values for all predicted AF2-predicted bicycle protein models.

(A) Unrooted phylogeny of the species whose genomes were used to identify *bicycle* genes used in this study. Phylogeny branch lengths are proportional to the number of

substitutions per site in all concatenated conserved proteins (17) and the scale bar represents 0.1 substitutions per site. Color squares are the same colors used in panel (C) and (E).
(B) *Bicycle* gene expression in different morphs of *Schlechtendalia chinensis*. Morph symbols as follows: AM = autumn migrant from primary host, *Rhus chinensis*, to secondary host *Plagiomnium maximoviczii*; FM = Sexual Female; FU = Fundatrix; GL = Fundatriginae; L1 = first-instar nymph on secondary host; L2 = second-instar nymph on secondary host; MA = sexual male; MF = sexual female; SE = spring migrant to primary host.
(C) Distribution of pLDDT for bicycle proteins by aphid species.
(D) pLDDT values versus protein length colored by density.
(E) The same data shown in (B) colored by aphid species.
(F) pLDDT values versus MSA alignment size used for AF2 predictions.

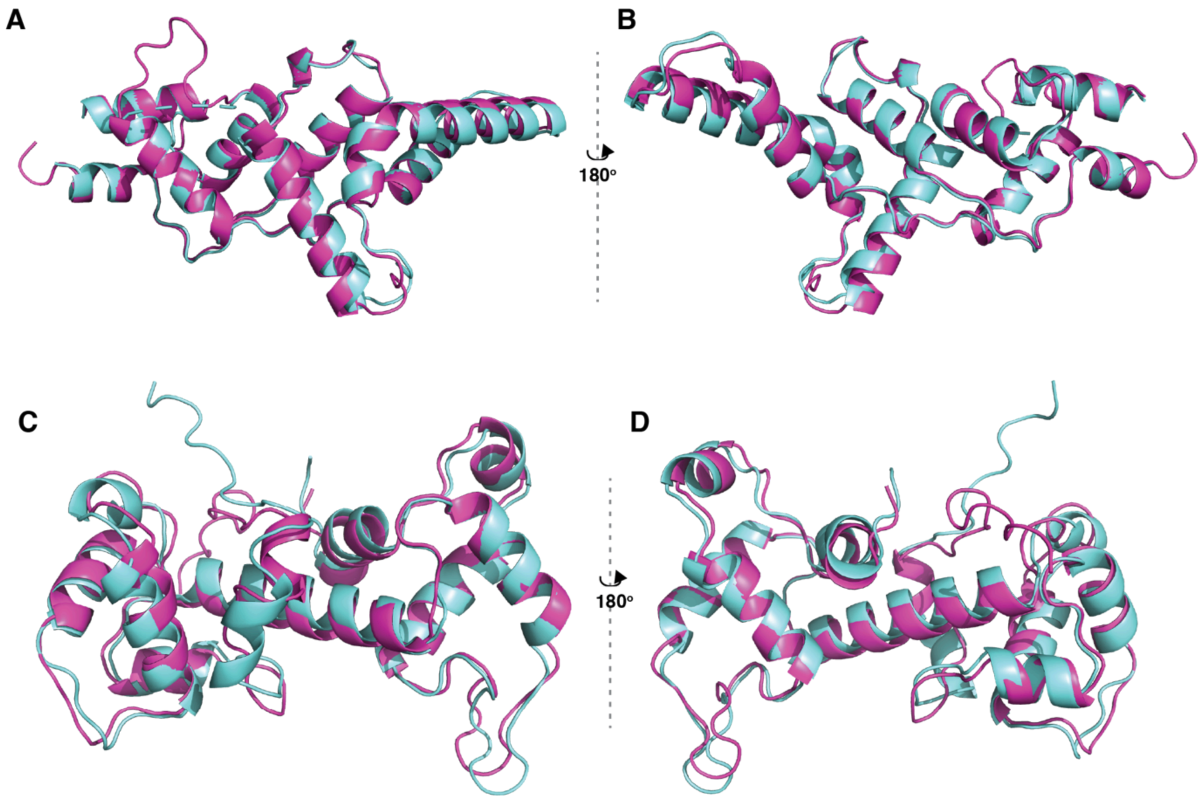

**Figure S12**: AF2-predicted models (cyan) of (A, B) g3873 and (C, D) g2703 using custom MSAs superimposed on their respective crystal structures (magenta).

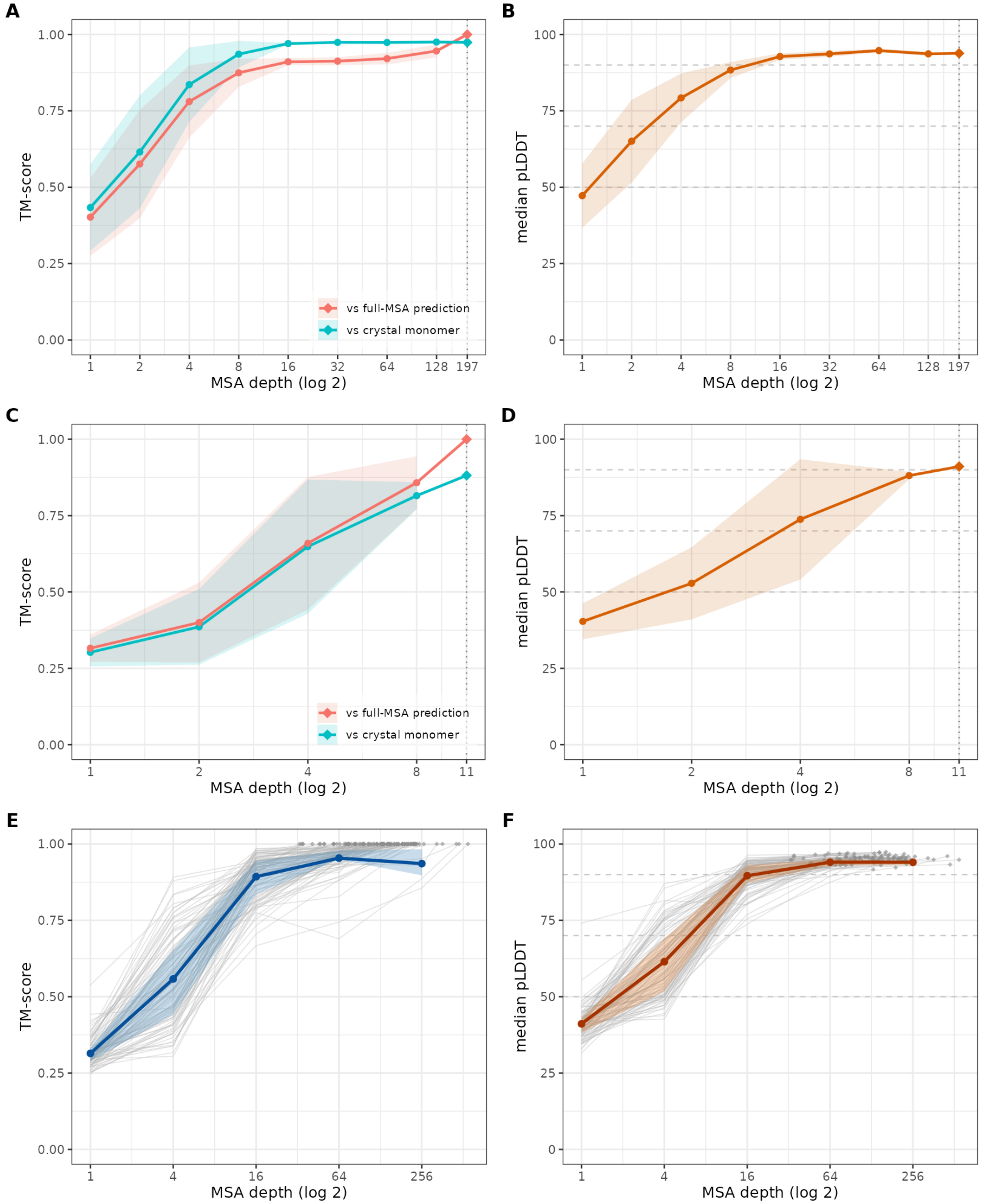

**Figure S13:** Replicated downsampling of multiple sequence alignments (MSA) reveals minimum MSA depth required for high accuracy and high confidence protein structure predictions with AF2.

(A-D) AF2 predictions of G3873 (A, B) and G2703 (C, D) reveal that similarity, measured as TM-score to the crystal structures and to the full-MSA predictions (cyan and red, respectively, in A and C) increases with greater MSA depth up to about 16 homologs. A similar pattern is observed for median pLDDT of each prediction (B and D).

(E-F) Similar patterns are observed for 100 randomly selected Bicycle proteins that were predicted with high confidence by AF2 with their full MSA. Prediction accuracy compared to the prediction with the full MSA (TM-score) and prediction confidence (pLDDT) plateau above MSAs with at least 16 homologs.

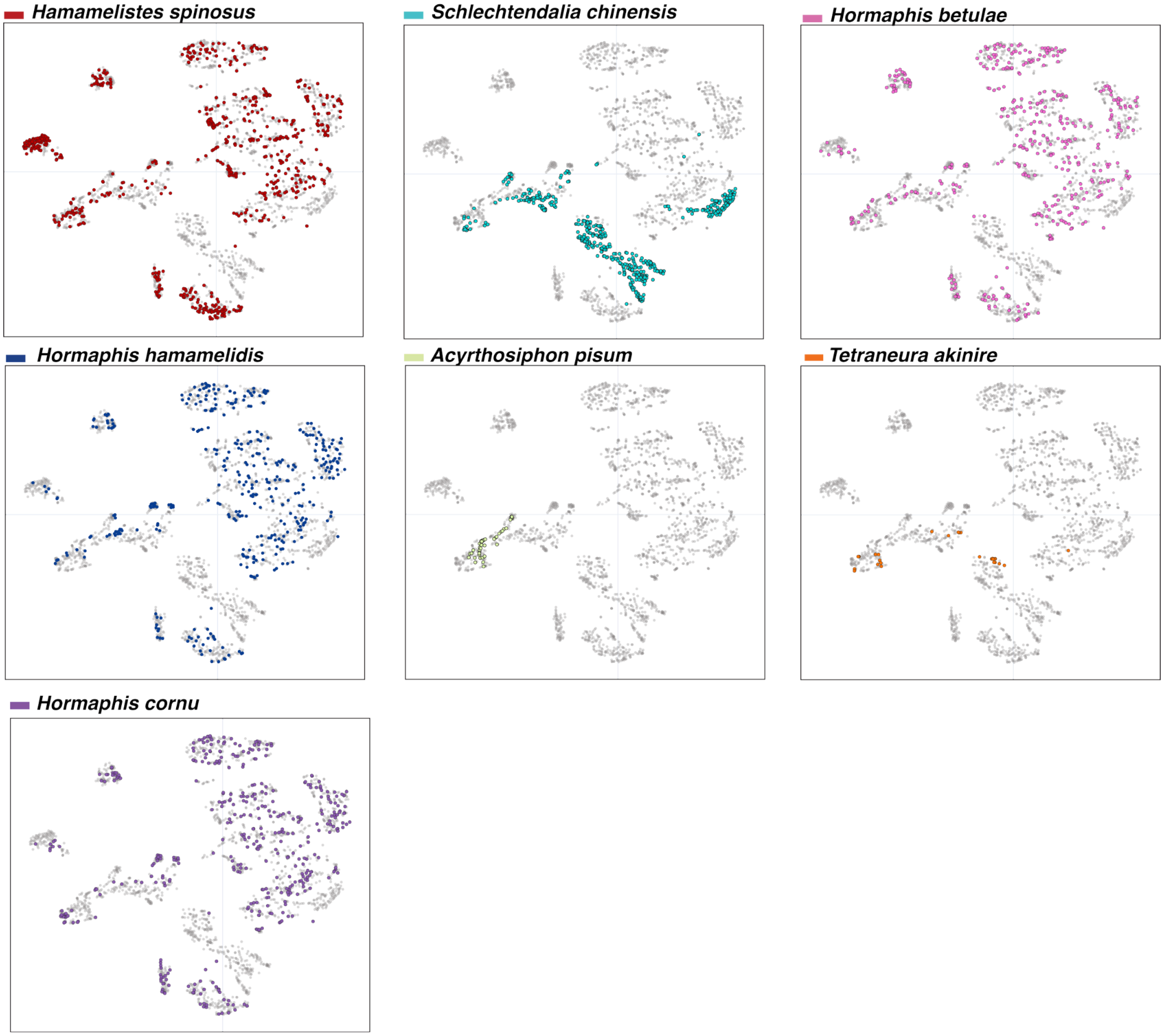

**Figure S14:** The t-SNE plot of AF2-predicted bicycle protein structures for seven aphid species from Figure 4B, with the species of each protein colored separately in each panel for each of the seven species.

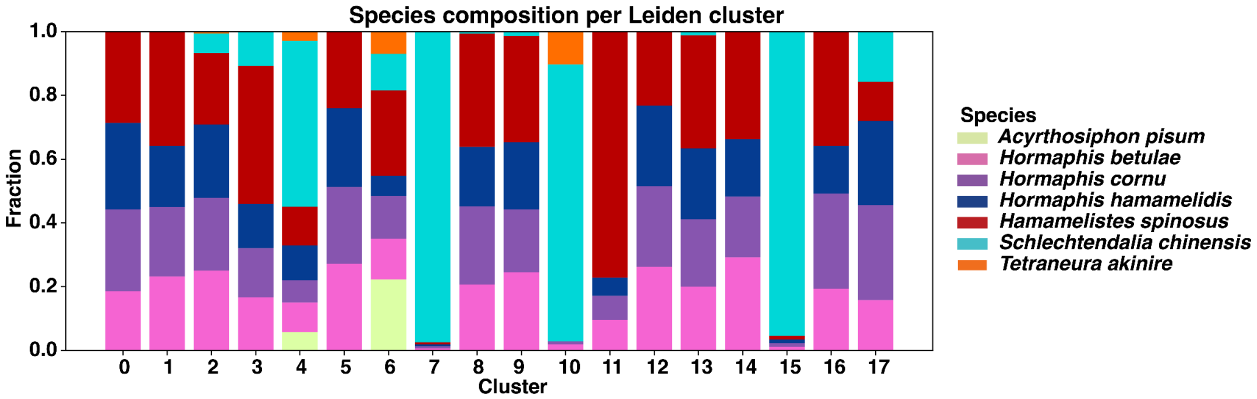

**Figure S15:** Bar graphs depicting the proportional contribution of proteins from each species to each Leiden cluster in the t-SNE plot from Figure 4B.

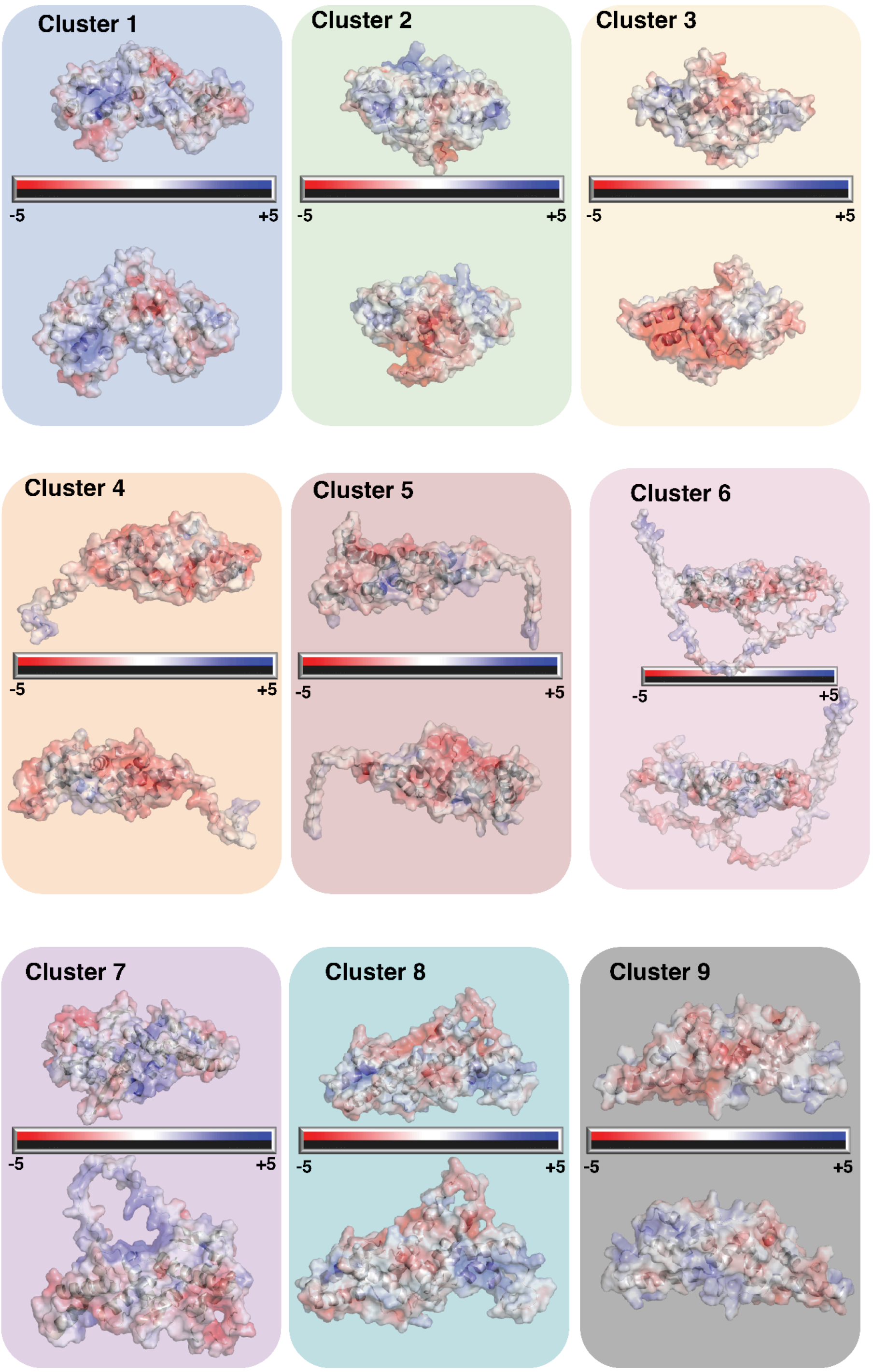

**Figure S16:** Space filling representation (two views) of APBS electrostatic surface potentials for Leiden cluster medoid models from Figure 5. Electrostatic potentials were computed for the nine cluster medoids using pdb2pqr (AMBER, pH 7.0) and APBS (linearized Poisson–Boltzmann, $\varepsilon_{prot} = 2$, $\varepsilon_{solv} = 78$, 0.15 M monovalent ions). Potentials were mapped onto the molecular surfaces in PyMOL using a fixed color ramp of −5 to +5 kT/e (white ≈ 0).

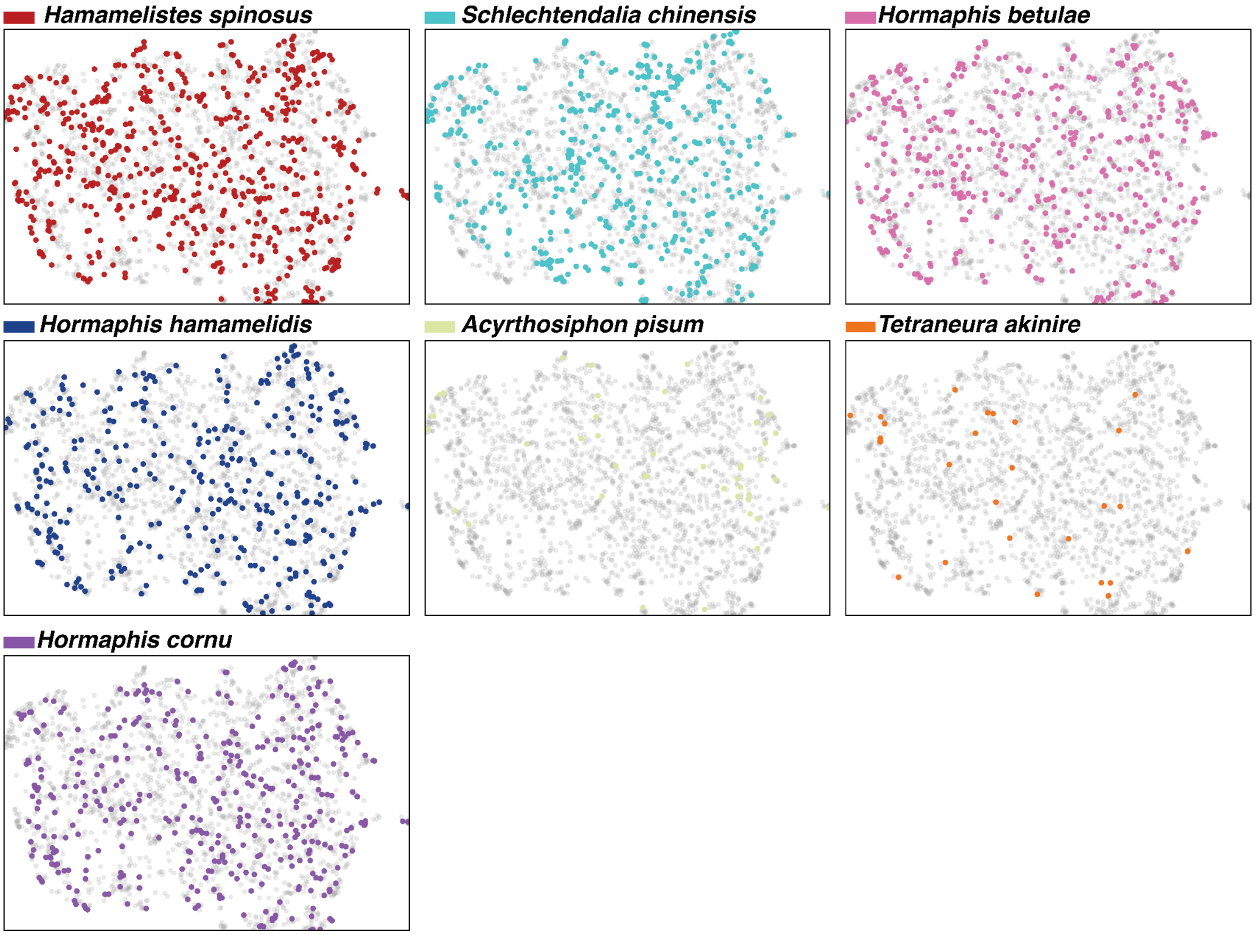

**Figure S17:** UMAP representation from Figure 5, with the species of each protein colored separately in each panel for each of the seven species.

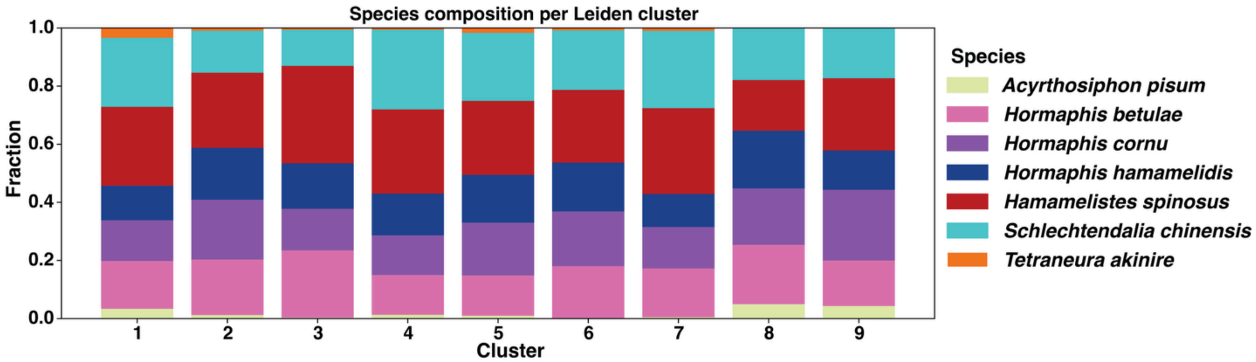

**Figure S18:** Bar graphs depicting the proportional contribution of proteins from each species to each Leiden cluster in the UMAP plot from Figure 5.

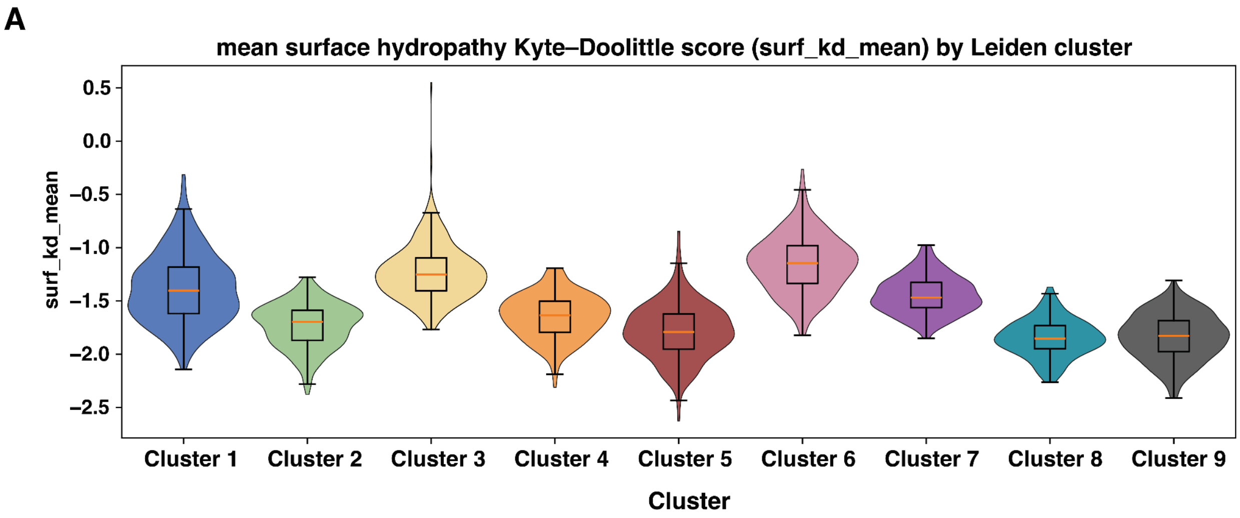

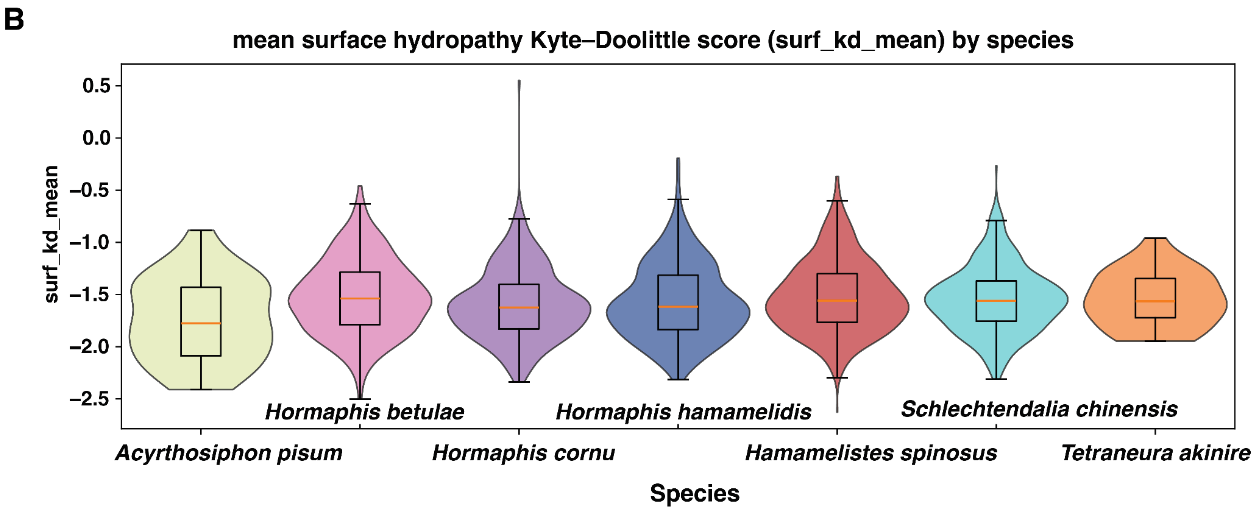

**Figure S19:** Violin plots of physicochemical property surface hydropathy measured using Kyte-Doolittle score weighted over the solvent exposed surface area by cluster (A) and by species (B).

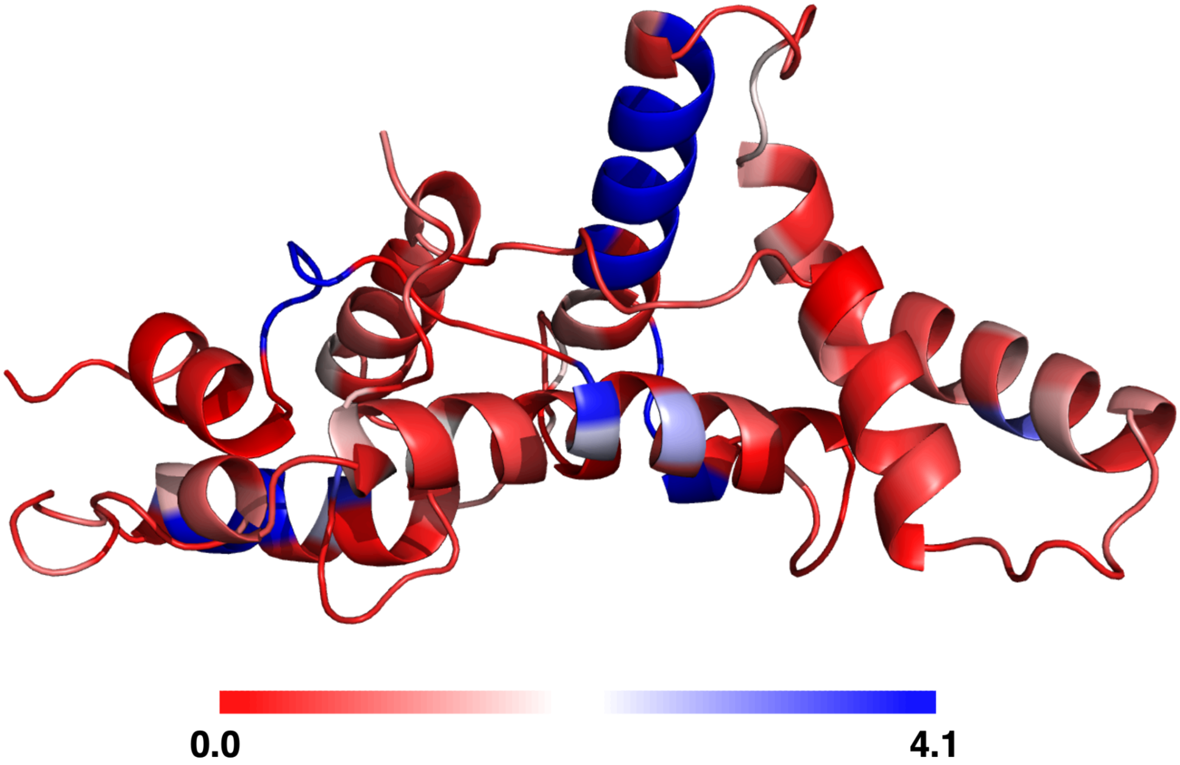

**Figure S20:** Per-residue sequence variability calculated as Shannon entropy (bits) mapped onto the backbone of the crystal structures of g3873. The ribbon is colored from blue (low entropy = conserved) to red (high entropy = variable) using a fixed scale (0–4.12 bits).

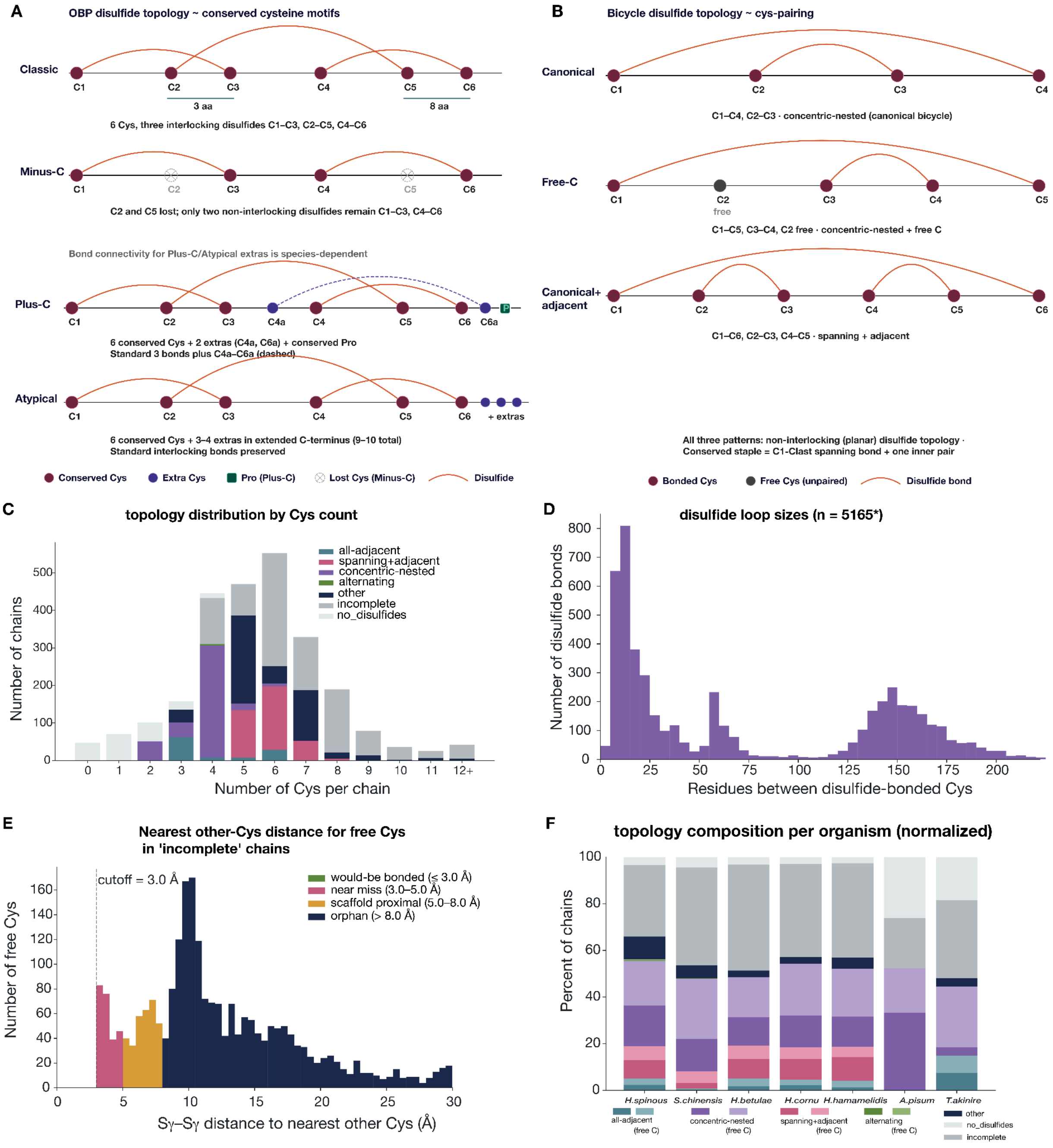

**Figure S21**: Disulfide topology schematics for OBPs and *H. cornu* predicted bicycle proteins.

(A) Topologies for Odorant binding protein (OBP) Disulfide topology schematic for four OBP classes: Classic, three interlocking bonds (C1–C3, C2–C5, C4–C6); Minus-C, loss of C2/C5; Plus-C, two extra Cys with a species-dependent C4a–C6a bond (dashed); Atypical, classic pattern plus 3–4 C-terminal Cys. The conserved Cys (red), extra Cys (blue), conserved Pro (green), lost Cys (open crossed circles), disulfides (orange arcs).

(B) Topologies for Bicycle disulfide topology for H. cornu predicted bicycle proteins models. Canonical (C1–C4, C2–C3), Free-C (C1–C5, C3–C4, C2 unpaired; grey), and Canonical + adjacent (C1–C6, C2–C3, C4–C5). All share a non-interlocking planar topology with a C1–C(last) staple plus one inner pair.

(C) Topology distribution by cysteine residue count for Bicycle proteins. Predicted models binned by Cys per chain (0–12+), colored by topology class. The distribution peaks at 6 Cys; concentric-nested and spanning + adjacent dominate at 4–6 Cys, while models with ≥8 Cys are largely incomplete.

(D) Disulfide loop sizes for predicted Bicycle proteins (n = 5,165). Bimodal distribution of residues between bonded Cys, with a short-range mode at ~10–20 residues corresponding to the first saposin-like half and a broad long-range mode at ~145–160 corresponding to the helix-swapped saposin-half. (E) Nearest available Cys distance to identify possible disulfide bonds in the incomplete group. Sγ–Sγ distance classified as would-be bonded (≤ 3.0 Å), near miss (3.0–5.0 Å), scaffold proximal (5.0–8.0 Å), or orphan (> 8.0 Å). Most free Cys are orphans, peaking near ~10 Å.

(F) Common disulfide connectivity patterns in predicted Bicycle proteins. Cys-pairings are shown as normalized bar charts per species. The 1–4,2–3 concentric-nested bicycle is most common, followed by the three-bond spanning-adjacent topology.

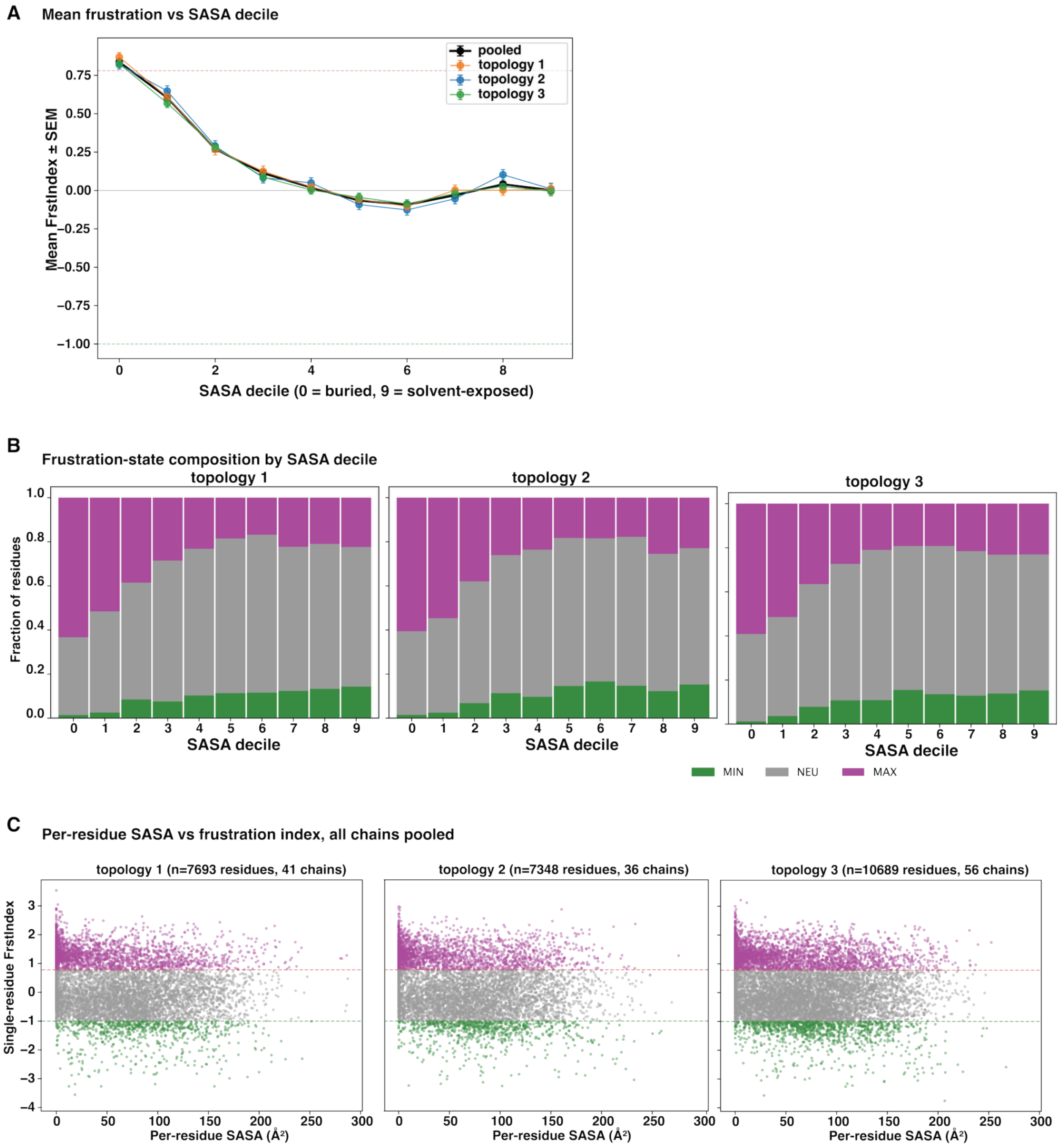

**Figure S22:** Frustration analysis for a subset of H. cornu bicycle proteins. (A) Mean frustration vs. SASA decile. Residues binned into SASA deciles (0 = buried, 9 = solvent-exposed); mean single-residue FrstIndex ± SEM shown for the pooled set and for each topology (1–3). Dashed lines mark the FrustratometeR MAX (+0.78) and MIN (−1.0) thresholds. All three topologies show the canonical buried-residue frustration excess, decaying to near-neutral or slightly minimally frustrated values by decile 4.

(B) Frustration-state composition by SASA decile. Stacked fractions of MIN (green), NEU (grey), and MAX (magenta) residues per SASA decile, split by topology. The buried deciles (0-1) are dominated by MAX-frustrated residues across all three subgroups; minimally frustrated residues accumulate toward the exposed deciles.

(C) Per-residue SASA vs. frustration index, all predicted models pooled. Scatter of single-residue FrstIndex against per-residue SASA ($Å^2$) for each topology (n / chains indicated). Points are colored by frustration state using the MAX (+0.78, magenta dashed) and MIN (−1.0, green dashed) cutoffs. Highly frustrated residues cluster at low SASA (<50 $Å^2$), while minimally frustrated residues are distributed across the SASA range.

**Table S1**: Bicycle proteins tested for recombinant expression.

| bicycle protein | *E. coli* expression | *SF9* cell expression | Crystals |
|---|---|---|---|
| g3191 | No | Yes | Yes |
| g3873 | Yes | Yes | Yes |
| g7134 | No | Yes | No |
| g15183 | No | Yes | No |
| g16073 (dgc) | Yes | Yes | No |
| g2703 | Yes | Not tested | Yes |
| g2079 | No | Not tested | Not tested |
| g2080 | Yes | Not tested | Not tested |
| g3028 | No | Not tested | Not tested |
| g3218 | No | Not tested | Not tested |
| g6303 | No | Not tested | Not tested |
| g10552 | No | Not tested | Not tested |
| g11197 | No | Not tested | Not tested |
| g11198 | No | Not tested | Not tested |
| g11199 | No | Not tested | Not tested |
| g13344 | No | Not tested | Not tested |
| g16009 | No | Not tested | Not tested |
| g16059 | No | Not tested | Not tested |

**Table S2**: Crystal data collection parameters and structure statistics for g2703 and g3873.

| | Se-Meth | Se-Meth | Sulfur-sad | Native |
|---|---|---|---|---|
| | High-Remote | Low-Remote | | |
| Protein name | g2703 | | g3873 | |
| PDB ID | 9ZML | | 12AC | |
| Data Collection Beamline | SSRL-12-1 | SSRL-12-1 | SSRL-12-1 | SSRL-12-1 |
| Space group | P 21 21 21 | P 21 21 21 | P21 21 21 | P21 21 21 |
| Wavelength | 0.964 | 0.984 | 1.573 | 0.978 |
| Cell dimensions | | | | |
| a, b, c (Å) | 40.986, 75.996, 85.808 | 41.002,76.012, 85.827 | 66.12, 71.268, 193.433 | 65.98, 71.17, 193.077 |
| alpha, beta, gamma (°) | 90.00, 90.00, 90.00 | 90.00, 90.00, 90.00 | 90.00, 90.00, 90.00 | 90.00, 90.00, 90.00 |
| Number of unique reflections | 53,411(5,195) | 53148(4881) | 16045(974) | 53700(3568) |
| Resolution range (Å) | 38-1.4 | 38-1.4 | 40-3.1 | 38.9-2.1 |
| Rsym | 0.023(0.84) | 0.024(0.91) | 0.031(0.873) | 0.021(0.781) |
| I / σI | 10.8(0.89) | 10.9(0.92) | 12.7(1.57) | 5.5(1.27) |
| $CC_{1/2}$ | 0.999(0.273) | 0.999(0.269) | 0.99(0.371) | 0.997(0.291) |
| Completeness (%) | 99.7(98.52) | 99.12(92.8) | 90.04(94.2) | 99.45(98.4) |
| Redundancy | 53.2(43.9) | 52.1(36.06) | 41.5(39.4) | 6.7(6.4) |
| Refinement | | | | |
| Resolution (Å) | 37.99-1.40 | | | 38.96-2.10 |
| Rwork / Rfree | 0.1866/0.2053 | | | 0.2275/0.2555 |
| No. atoms | | | | |
| Protein | 1452 | | | 5374 |
| Ligand/ion | 6 | | | - |
| Water | 271 | | | 229 |
| B-factors | | | | |
| Protein | 30.41 | | | 56.82 |
| Ligand/ion | 51.3 | | | - |
| Water | 43.01 | | | 51.34 |
| R.m.s. deviations | | | | |
| Bond lengths (Å) | 0.005 | | | 0.009 |
| Bond angles (°) | 0.835 | | | 1.094 |

**Table S3:** Disulfide bond parameters in the X-ray structure of G3873.

| Chain | Motif | Res 1 | Res2 | Chi1 ($\chi$ 1) | Chi2 ($\chi$ 2) | Chi3 ($\chi$ 3) | Distance (Å) | Chi2' ($\chi$ 2') | Chi1' ($\chi$ 1') | Disulfide Strain Energy (kJ/mol) |
|---|---|---|---|---|---|---|---|---|---|---|
| A | N-terminal Saposin-like (CYC) | 30 | 191 | 179.89 | -89.01 | -70.41 | 2.04 | -67.43 | -61.91 | 7.971993 |
| A | non Saposin-like | 38 | 48 | -71.09 | -136.58 | 98.06 | 2.03 | -64.94 | -75.7 | 15.178506 |
| A | C-terminal Saposin-like (CYC) | 92 | 120 | -74.99 | -61.82 | -77.47 | 2.09 | -84.16 | 173.03 | 8.296798 |
| B | N-terminal Saposin-like (CYC) | 30 | 191 | -176.11 | -97.13 | -76.26 | 2.05 | -74.56 | -62.46 | 9.609778 |
| B | non Saposin-like | 38 | 48 | -70.7 | -134.93 | 97.59 | 2.02 | -68.08 | -73.45 | 14.779842 |
| B | C-terminal Saposin-like (CYC) | 92 | 120 | -74.11 | -61.7 | -79.96 | 2.06 | -84.98 | 171.34 | 8.278995 |
| C | non Saposin-like | 38 | 48 | -55.56 | -85.59 | -40.94 | 2.03 | 90.74 | -160.81 | 29.527321 |
| C | C-terminal Saposin-like (CYC) | 92 | 120 | -71.07 | -63.7 | -75.4 | 2.07 | -83.87 | 170.17 | 8.024059 |
| D | N-terminal Saposin-like (CYC) | 30 | 191 | -174.79 | -88.35 | -77.99 | 2.05 | -87.85 | -60.28 | 10.140293 |
| D | non Saposin-like | 38 | 48 | -83.74 | -135.05 | 107.05 | 2.04 | -77.57 | -80.6 | 25.848478 |
| D | C-terminal Saposin-like (CYC) | 92 | 120 | -74.94 | -61.97 | -78.27 | 2.08 | -84.62 | 173.58 | 8.218305 |

**Table S4**: Foldseek hits for g3873 and g2703 identify only poor quality matches (.xlsx file).

**Table S5:** TM-scores for all saposin-like proteins compared to CYC-domains in g3873 (.xlsx file).

| | | | | |
|---|---|---|---|---|
| RMSD (Cα matched) | structure | $RMSD = \sqrt{\left(\frac{(1)}{(N)}\right)\sum_{i=1}^{N} d_i^2}$ | Å | Higher = less similar; lower RMSD = closer match |

**Table S6:** TM-scores for all saposin-like proteins compared to the sapsin-like CYC-domains in g2703 (.xlsx file).

**Table S7:** Filtered data of AF2-predicted models for all seven aphid species (.xlsx file).

**Table S8:** List of physicochemical properties examined for all high-confidence AF2-predicted bicycle proteins.

| Column | From | Property | Units | Interpretation |
|---|---|---|---|---|
| pdb_file | Metadata | ID | — | Identifier |
| species | Metadata | Species label | — | Identifier |
| status | Sanity check | QC flag | — | Pass/fail/category |
| seq_len | Sequence | Length | residues | Larger protein |
| seq_frac_pos | Sequence | Frac. {K, R, H} | 0–1 | More basic residues |
| seq_frac_neg | Sequence | Frac. {D, E} | 0–1 | More acidic residues |
| seq_frac_hydrophobic_FLIV | Sequence | Frac. {F, L, I, V} | 0–1 | More hydrophobic composition |
| seq_net_charge | Sequence | n(pos) − n(neg) | net residues | More net positive sequence |
| sasa_total | Structure | Total SASA | Å$^2$ | More exposed surface |

| | | | | |
|---|---|---|---|---|
| sasa_hydrophobic | Structure | SASA on Hydrophobic set | Å$^2$ | More hydrophobic surface exposure |
| sasa_pos | Structure | SASA on Positive set | Å$^2$ | More exposed positive surface |
| sasa_neg | Structure | SASA on Negative set | Å$^2$ | More exposed negative surface |
| frac_sasa_hydrophobic[1] | Structure | Hydrophobic surface fraction | 0–1 | Surface more hydrophobic overall |
| surf_kd_mean[2] | Sequence and structure | Weighted KD mean (exposed) | KD (unitless) | Exposed surface more hydrophobic on avg |
| surf_kd_median[3] | Sequence and structure | KD median (exposed) | KD (unitless) | Typical exposed residue more hydrophobic |
| roughness_sasa_over_hull[4] | Structure | Roughness proxy | unitless | More corrugated/indented surface |
| hydropatch_count | Structure | No. of hydrophobic patches | count | More fragmented hydrophobic surface |
| hydropatch_mean_size[5] | Structure | Mean patch size (Σ SASA_res) | Å$^2$ | Larger typical hydrophobic patches |
| hydropatch_max_size[5] | Structure | Max patch size (Σ SASA_res) | Å$^2$ | One dominant “sticky” patch |
| surface_charge_dipole[6] | Structure | Charge dipole magnitude \|D\| | scaled charge·Å | Stronger +/− polarization |
| exposed_net_charge[7] | Structure | Net exposed charge | net residues | More net positive exposed charge |
| exposed_net_charge_over_sasa[8] | Structure | Exposed charge density | residues/Å$^2$ | Higher surface charge density |
| net_charge_over_sasa[8] | Sequence and structure | Seq charge density | residues/Å$^2$ | Higher sequence net charge per area |
| amphipathic_helix_fraction[9] | Sequence and structure | Amphipathic helix fraction | 0–1 | More amphipathic helix character |
| max_hydrophobic_moment[10] | Sequence and structure | Max hydrophobic moment | Unitless | Stronger amphipathic helices |

**Details:**

- Residue sets: Hydrophobic {I, V, L, F, C, M, A, W, Y, P}; Positive {K, R, H partial at pH 7.0}; Negative {D, E}.
- Exposure threshold θ = 5 Å$^2$.

- Dipole scaling $\alpha = 1/100$.
- Hydrophobic patches: DBSCAN on exposed hydrophobic sidechain centroids (eps = 7 Å, min_samples = 3), patch size = Σ SASA_res

1. **Fraction hydrophobic SASA:**

$$frac_{sasa_{hydrophobic}} = \frac{\left(SASA_{hydrophobic}\right)}{\left(SASA_{total}\right)}$$

2. **SASA-weighted surface KD mean (exposed):**

$$KD_{mean} = \frac{\left(\Sigma_{exposed} KD(res)\cdot SASA_{res}\right)}{\left(\Sigma_{exposed} SASA_{res}\right),\ \ SASA_{res} \geq \theta}$$

3. **Surface KD median (exposed):**

$$KD_{median} = median\{ KD(res)\} over\ exposed\ residues, \quad SASA_{res} \geq \theta$$

4. **Roughness proxy:**

$$roughness = \frac{SASA_{total}}{A_{hull}}$$

5. **Hydrophobic patch size (cluster j):**

$$size_j = \Sigma_{\{res \in cluster\ j\} SASA_{res}}$$

6. **Surface charge dipole magnitude:**

$$D = \Sigma_{exposed} q(res)\cdot \left(SASA_{res} \cdot \alpha\right)\cdot r_{centroid(res)}; \quad |D| = \sqrt{Dx^2 + Dy^2 + Dz^2}$$

7. **Exposed net charge:**

$$exposed_{net_{charge}} = \Sigma_{exposed} q(res)$$

8. **Charge densities:**

$$exposed_{net_{charge_{over_{sasa}}}} = \frac{exposed_{net_{charge}}}{SASA_{total}};$$

$$net_{charge_{over_{sasa}}} = \frac{seq_{net_{charge}}}{SASA_{total}}$$

9. **Amphipathic helical fraction:** $\frac{N_{covered}}{L}$

where $N_{covered}$ is the number of residues covered by at least one helical window $\mu_H(i) \geq 0.35$, and $L$ is sequence length, helical mask $h_j = 1$ from $\phi \in [-100^{\circ}, -30^{\circ}]$, $\psi \in [-80^{\circ}, -5^{\circ}]$ plus min helix run length ≥ 7.

10. **Max hydrophobic moment:**

$$\mu_{H(i)} = \frac{1}{W} \sqrt{\sum_{k=0}^{W-1} H_{i+k} \cos cos\,(k\delta))^2 + (\sum_{k=0}^{W-1} H_{i+k})^2}$$

$$max_{hydrophobic_{moment}} = max_{i:\, h_{i:i+W-1}=1}\, \mu_{H(i)}$$

**Table S9:** Physicochemical properties of all 2400 bicycle proteins (.csv file)

**Table S10:** Metric used to generate the overall structure space of proteins

| Metric | Derived from | Formula / Definition | Units | Interpretation |
|---|---|---|---|---|
| TM-score | structure | $TM = \frac{\left(\frac{1}{L_{norm}}\right) sum_{i=1}^{N} 1}{1 + \left(\frac{d_i}{d_0}\right)^2}$ <br> $d_0 = 1.24 * \left(L_{norm} - 15\right)^{\frac{1}{3}} - 1.8\ \text{Å}$ | unitless (0–1-ish) | Higher = more similar global fold (less sensitive to local outliers than RMSD) |

**Table S11:** Data and statistics of newly assembled and re-annotated genomes (.xlsx file).
